# A topological connectome surface network with spin-half particles can produce brain-like signals and store memory

**DOI:** 10.1101/2022.08.01.502331

**Authors:** Siddhartha Sen, Tomas J. Ryan, David Muldowney, Plamen Stemanov, Maurizio Pezzoli

## Abstract

In this paper we use the methods of theoretical physics to show how brain-like signals can be generated in a special surface network with the topological connectivity of the biological brain by exploiting its form. The network is required to have surface spin-half particles. We show that the signals thus generated carry information regarding their creation and can transfer and store this information to form memory structures in helical aligned surface spin-half particles present on the surfaces of the pathways traversed by the signals. Theoretical neuroscience is progressing strongly with novel representations of the brain, enhanced by the increase of computational power now available. New methods to explore complex brain events and the structures for storing memories as engrams are emerging. However, there are major conceptual theoretical problems that remain unaddressed. Current theoretical methods are very capable of reacting to experimental results and modelling both neural signalling and structure. Yet they still fall short to throw light on how the brain creates its own information code, or relate the variety of brain signals observed, or explain where and how memories are stored. We prove that current brain signal interpretations cannot carry information regarding their formation so that they cannot be used to understand how memories of events are related to signals. Thus, our results address these basic unresolved theoretical problems of neuroscience and suggest testable solutions. The memory structures of aligned spin-half particles suggested have not been observed in biological organisms as yet they but they have been observed in solid state physics and their existence is consistent with conventional understandings of neurobiology. All the results stated follow from the dynamical law for the network.

## 1 Introduction

Experimental progress in neuroscience has been remarkable. New tools and methods of studying the brain have led to insights regarding neural activity and memory storage. The consensus is that sensory input signals carry information that processed by the brain, gives rise to our sensory experiences. By internal mechanisms the brain evaluates saliency and relevancy of the sensory experience, which might trigger the storage of the event within the neural circuit as long-term memory. The process of memory creation has been studied in depth [1] but where memories are actually stored is still under investigation [2]. It is generally accepted that memory formation is linked to information carried by propagating action potentials that traverse brain circuits. The action potential generation is well described by the Hodgkin Huxley equations [3]. They are created as a result of inflows and outflows of ions controlled by protein channels present in neural membrane and their properties. The process is initiated when action potentials enter the soma, the cell body of a neuron, through its signal receiving dendrites. If the input signals cross a certain threshold value, a processed action potential exits the neuron via the one exit neuron fibre of the soma, its axon. The signals are observed electrical voltage pulses and are assumed to satisfy Ohm’s law of electromagnetism. They are dissipative waves.

On the theoretical side considerable progress in modelling assembly of neurons has been made that are capable of reacting to experimental results [4, 5] and even predicting the possibility of unexpected brain excitations [6] but there is a fundamental problem in modelling signal production that needs to be rectified. The problem is that all exiting signal generating models such as the Hodgkin Huxley model or its variants [3] create dissipative waves that are not capable of carrying information. This will be proved and is our first unexpected result. For dissipative waves there is a causal cycle of energy injection and energy dissipation. For existing ion flow models the energy injection comes from membrane ion inflows and outflows using special membrane ion channels gates, that lead to a potential gradient that provides energy, which is dissipated by the ohmic current created. This causal cycle of energy injection and dissipation propagates the signal. It is driven by the membrane properties, the active element of the process, and does not depend on how the signal was started, which means that the individual action potential pulses cannot carry information regarding the input signal that created them [3].

But we do know that observationally action potentials are produced directly in response to excitatory inputs and we also do know that the information carried by action potentials evoked by our senses convey messages to the brain that allows us to see, taste, hear, and feel sensations. This is a fundamental problem that needs rectification.

This theoretical gap is currently addressed in neuroscience [7] by suggesting that information from sense organs is carried, not the pulses themselves, but by their production rate and/or by the way they are distributed. But how this is done is not theoretically known. Such a situation is unsatisfactory as signal pulses are generated by inputs from sensory or other organs and a proper understanding of how this information is transferred to signals is essential if we want to link signals and memory in a clear way.

There is another non-ion flow model available for generating propagating brain signals [8]. Where it is shown theoretically, and subsequently experimentally confirmed that brain membrane distortions can produce non-dissipative acoustic pulses called solitons. Subsequently it was shown that these acoustic solitons induce voltage spikes similar to those observed for action potentials [9]. Perhaps these non-dissipative excitations can carry information regarding their formation? Unfortunately, this is not the case. The process of generating acoustic solitons depends in an essential way on an underlying thermodynamic phase transitions process. But thermodynamic processes described by thermodynamic state functions [10], have no memory of how they are formed. Thus, the acoustic solitons idea does not resolve the theoretical problem.

To place these problems in context we recall that there is compelling experimental evidence that a complex chain of biological processes take place within neurons during memory creation, such as long-term and short-term synaptic plasticity [1], but less is known about how memories (and the information within them) are stored. What is known is that memories are not stored in independent neurons or their synapses [11], but perhaps non locally in an unknown way, in the pathways between special group of neurons, in a structure called an engram [12, 13, 14]. There is experimental support for this picture [15], but the lack of a theory for engrams is problematic [2]. Ideally, a theory should explain how action potentials carry information, if not in the pulses themselves, then in their production rates or distributions in a way that is theoretically understood. The theory should then be able to explain how such information is transferred from the signal and stored as memory. Each step should be testable. Currently there is no such explanation available.

A related basic unanswered question is to suggest how information stored in the brain is retrieved. The list of problems stated are all fundamental questions that need to be addressed taking note that current observational neuroscience memory research reveals [14] that global features of the brain are important for memory and for its functioning as a distributive global system.

### 1.1 Outline of results

In this paper we will suggest testable solutions to these fundamental problem by constructing a special surface network with surface spin-half particles and then show, using the methods of theoretical physics and mathematical physics, that it has the properties we desire.

Brain-like signals are generated in the network using an unconventional global method that involves the form of the network. A universal class of input signals for generating all signals is suggested and the response of the network to them is determined by a dynamical law. The universal input signals introduced change the connectivity of the network, and the global dynamical law requires that the mathematical structural properties of the network should remain intact during these topological changes. This rather abstract almost philosophical law will be progressively refined so that at the end it becomes an operational law that leads to the result that a wide variety of brain-like signals can be generated in the network by topology changing input signals and that the signals are solutions of the one-dimensional non-linear Schroedinger differential equation. We will explain the hidden enabling role of the surface spin-half particles of the network plays in signal generation.

We will show that the propagating signals carry information regarding their formation encoded in a self-generated manner. We will then show that there is a natural method for the signals to transfer the information they carry to form memories structures composed of aligned surface spin-half particles in the pathways they traverse. The memory structures have memory specific excitation frequencies that may be used to recall them. These frequencies are estimated. The results established are quantitative. Analytic expressions for brain excitations are available but the speeds and amplitude scales of these excitations depend on the choice of scale parameters for length, time and voltage. Thus, the memory structure that naturally emerges from the network is that of an engram [16].

The propagating signals of the network are one-dimensional non-dissipative solitons which we identify them as action potentials. We justify this identification by fitting observational data for action potentials to explicit soliton solutions available and by carrying out three calculations. In the first calculation we show that it is possible to estimate the membrane resting voltage by assuming the network surface has the electrical properties of the brain membrane [17] and a theoretical result [18] based on quantum electrodynamics. We then use this result and the laws of low Reynold number [19] physics to correctly predict the nature of the observed membrane distortion waves that accompany soliton pulses. A formula is derived that shows that the membrane distortion wave amplitudes are proportional to the amplitudes of the soliton pulses that generates them. With these two results in place, we next tackle an apparently unrelated problem.

It is known that the insulating myelin sheaths that cover axons have regularly spaced gaps in them known as the nodes of Ranvier [20]. We use the formula for the resting membrane potential and the formula relating membrane distortion waves and soliton pulse amplitudes to estimate the gaps between the nodes of Ranvier. The estimate makes use of the networks method of signal generation and memory storage thus the result further supports the identification of solitons with action potentials.

We relate the existence of the gaps between the nodes of Ranvier to the process of memory structure creation by signals. Signals lose energy as they move due to the fact that they produce magnetic fields that act on surface spin-half particles and align them to form memory structures. This process requires energy. We estimate the energy loss for this aligning process and find that the energy loss is ≈ 100% after signals travel one centimetre. Thus, the signals need boosting. We identify the conditions required to boost signals in the networks and find that these conditions are met at the nodes of Ranvier [20]. Thus, these three calculations fit together and support the method of signal production and the identification of soliton pulses as action potentials. The novel features of the approach are a new topological way to generate signals and a suggested way to create memory structures that is based on the laws of electromagnetism [21].

## 2 Overview of method

Our approach is based on methods of theoretical physics and results of mathematics. We start by explaining the choice of surface network and its key properties before describing the topological method of signal generation and how memories can be created by signals.

### 2.1 The network is a Riemann surface with spin structure

The network has two layers of topological structures both of which play an essential role in both signal production and memory storage. Topology [22] is a mathematical discipline in which two geometrical objects are regarded to be the same if they can be converted to each other by continuous deformations. Thus, the surface of a sphere and that of a cube are topologically the same. A remarkable mathematical theorem of topology [23] tells us that any compact surface, no matter how complex, is topologically equivalent to either a sphere surface within handles or an orderly array of linked doughnuts. We choose our surface network to be an orderly array of joined doughnut surfaces. It thus describes any topological compact surface.

Structures that are topologically different can be described by topological numbers that can separate them into different topological classes. Our surface network has two integer topological numbers. The first, an integer *g*, is called the genus of the surface tells us how many linked doughnuts are present in the network. A sphere surface fits in the theorem and has genus *g* = 0, a torus *g* = 1 (Fig. 1A).

**Figure 1.**
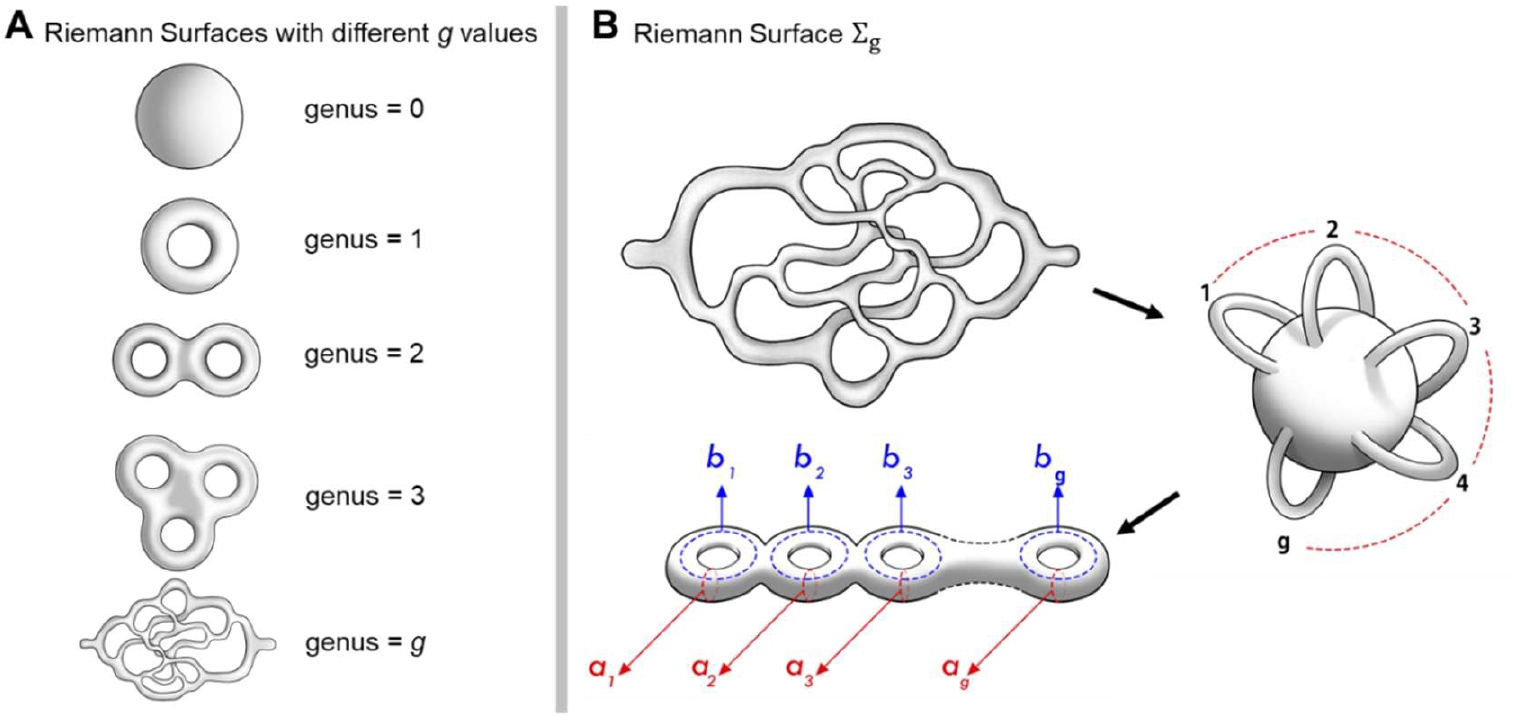
Topological approach to *Σ*_*g*_. **A.** Compact surfaces of 3D objects with increasing genus number. **B**. Topological connectivity property captured by loop coordinates round the surface of the *g* connected doughnuts [23]. There is a set of 2*g* of natural topological closed loop coordinates (*a*_*i*_, *b*_*i*_), *i* = 1, 2, …*g*.

The first topological number of the network thus gives information about its topological connectivity. The second topological number gives information about the topology of surface spin-half particles patterns on the network. Spin-half particles are magnetic and can thus link together to form topologically distinct patterns called spin structures. We will represent a spin structure by an integer *W* which will be define shortly.

What is the point of the doughnut construction? The answer is unexpected: the linked set of doughnut system is an exact surface representation of the topological connectivity of any brain connectome. Recall a connectome represents the connectivity architecture of a biological brain in which neurons are nodal points and axons are lines. If we replace each line of the connectome by a tube and each nodal point by a nodal region, we will create an enormously complex unknown surface in three dimensions. But the topology theorem we stated earlier tells us that this unknown complex surface is topologically equivalent to an orderly set of connected doughnuts. Currently computational neuroscientists are using the recently found connectome networks of a fly to use it to study the dynamical behaviour of the fly brain [24, 25]. Our surface network thus exactly captures an important topological property of the brain and has a mathematical description which makes it an interesting object to study (Fig. 1B).

At this stage we have an interesting surface network and we have made a claim that it can generate signals but to proceed we need a dynamical law and we need to introduce well defined input signals and explain how output response signals are generated. Our first statement of the dynamical law is that it requires that all dynamical activities of the network should preserve its fundamental structure. To operationalize the law, we need to enrich the network surface by adding more mathematical structure to it that will then define it.

Our first enrichment is to assume that the network surface is a smooth compact surface which has charges similar to the membrane cover of the brain’s assembly of three-dimensional individual neurons, and it has surface spin-half particles. The assumption of smoothness has profound consequence. It immediately turns our surface with surface spin-half particles, into a well-studied mathematical surface known as a Riemann surface with spin structure [26, 27, 28, 29].

### 2.2 Choosing a dynamical law

There are good reasons for selecting the abstract dynamical law stated for the network. It is because such a law will be valid for the brain. The brain is an open dissipative system that changes its structure over time. We have constructed a smooth surface representation of a brain connectome that captures its topological connectivity. Thus, our network has three key properties, it has topological connectivity, it has spin topology and it is smooth. But we know that the structure of the brain changes with time. New axons are created others are pruned and we know that these changes do not spoil the functioning of the brain. These changes modify the topology of the brain. Thus, any dynamical law proposed should have this property. The law should remain invariant under the topology changes of the brain described. Let us restate our dynamical law drawing attention to the fact it should be invariant under topology changes.

### 2.3 Dynamical law of network: First version

#### Any input surface disturbance and the response of the surface to it must respect the Riemann surface structure of the network

In essence the dynamical law requires the structural stability of the network. It means that even if new axons are created and others are pruned the topology can change but its Riemann surface nature should not change. The new axons or the pruning of axons are pruned or created can still preserve the Riemann surface structure of the network. Our signal generating or memory storage methods depend only on the network’s Riemann surface structure thus they will not be affected by topology changes. Surprisingly we will show that this rather non-detailed law with no ad-hoc parameters law is sufficient to show that brain excitations created by pinch deformations are solutions of the nonlinear Schroedinger equation.

We now start introducing the technical results and make some general comments. The first general comment is that to study the way a Riemann surface [28, 29] responds to input signals it is necessary to define a suitable function. This is the normal practice when modelling a system. For instance, to study the oscillations on the surface of a sphere spherical harmonic functions that depend on the variables that describe the sphere surface are introduced. In this case the appropriate function is a function called the Riemann theta function [28].

### 2.4 Universal input signal

The next crucial technical step is to fix the nature of input signals that can generate signals in the network. Here we are guided by a previous discovery [28]. By making local topology changing pinch deformations of a Riemann surface that reduces its topology to that of a sphere, generates one dimensional soliton pulses that are solutions of a non-linear differential equation. The nature of these deformations is shown (Fig. 2). This suggests that we introduce pinch deformations as the universal signal generating input signal of the surface network. But this step immediately requires that the dynamical law should be further enriched so that it can accommodate topology changes. Recall our general reasoning for selecting a dynamical law suggested that the law must be invariant under topology changes because biological changes of topology always occur in the brain, but now, motivated by this approach [28], such an invariance of the law is also required, as pinch deformations input signals change topology. A new idea, that topology changes can produce one dimensional pulsed signals, has emerged.

**Figure 2.**
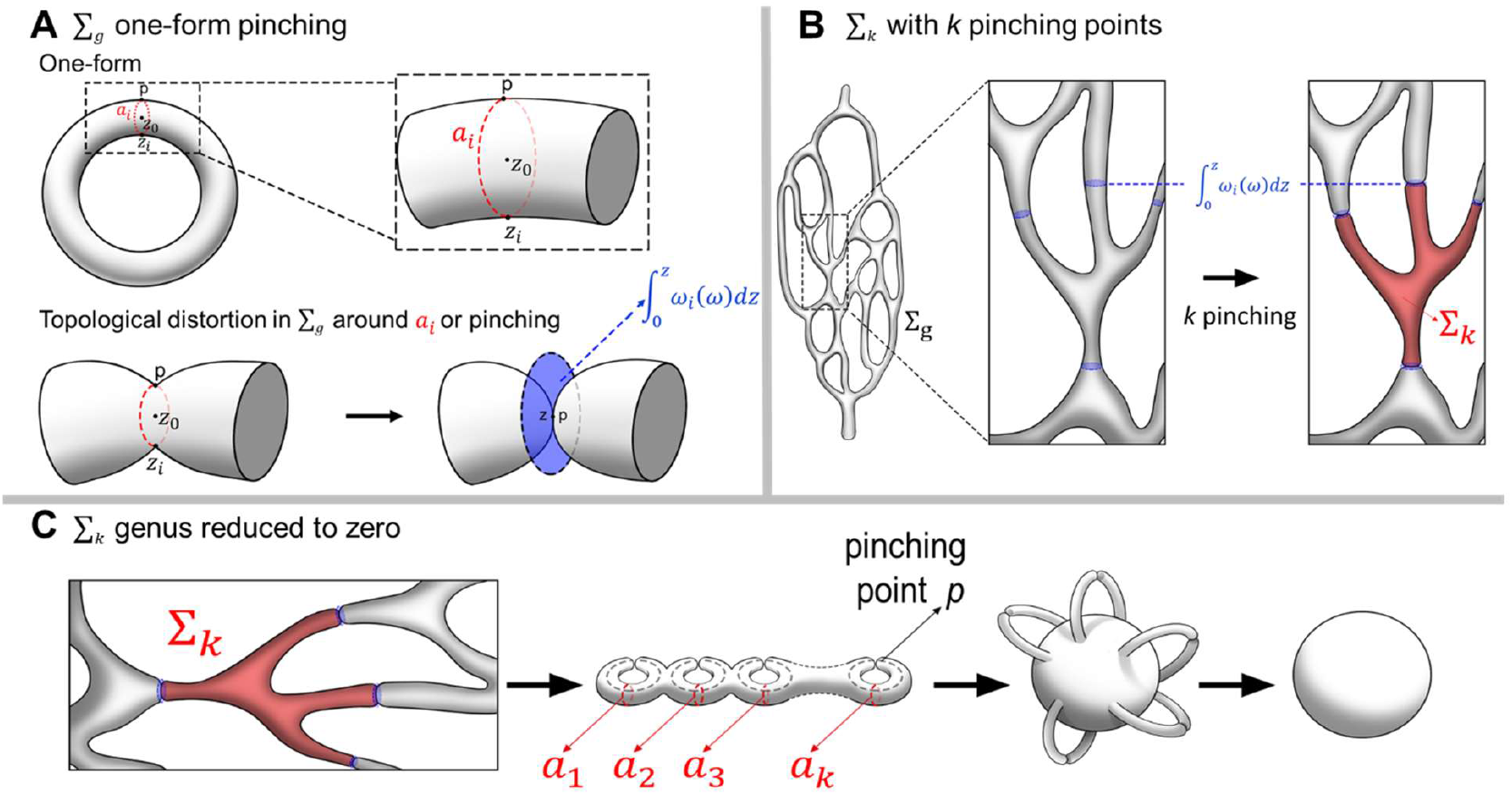
Local topological pinch deformations reduce, at a defined point, the circumference of a loop round a one-form *ω*_*i*_*(ω*) to zero. **A.** The dynamic law proposed is implemented by taking the limit of the two surface-points *p* and *z* approaching each other so that an arc joining them, a sub-section of an *a*_*i*_ loop connects them approaches zero. In this way the pinch deformation respects the Riemann Surface structure. This limit can be studied by considering the integral 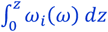 (see text for details). **B**. By pinching *k* one-forms that belong to Σ_*k*_, which is a sub-unit of the full surface-network Σ_*g*_. **C**. The subunit has its total genus value reduced to zero which transforms it transiently, into a topological sphere.

We next comment on the variables of the Riemann theta function. All the variables of the theta function are constructed from the variables of its associated Riemann surface. The theta function has variables belonging to two classes. It has *g* complex variables that emerge from one complex variable that represents a point on the Riemann surface, as we will show, and it has 2*g* discrete variables, called characteristics. At first these discrete variables seem to be unrelated to the variables of the Riemann surface, but we will explain later that they are related to the spin structure of the Riemann surface [26, 27].

We next comment on three technical issues. The first is to define pinch deformations in mathematical terms where it is clear that they change the topology of a signal producing subunit of *Σ*_*g*_ to that of a sphere. Second is to show that pinch deformations of *Σ*_*g*_ automatically lead to pinch deformed variables of Θ_*g*_. The third step is to explain that Θ_*g*_ has to satisfy a non-linear algebraic identity, the Fay trisecant identity if it is to represent a Riemann surface. This constraint is the key idea used to generate signals as pinch deformations convert the Fay identity for theta functions to a non-linear differential equation that the theta function, with pinch deformed variables, must now satisfy if it is to represent its associated Riemann surface.

There is a final twist. The deformed Fay identity leads to the non-linear Schroedinger equation only if *Σ*_*g*_ represents a special algebraic equation called the hyperelliptic equation [30, 31]. Thus, pinch deformations produce responses that are pinch deformed theta function solutions of the non-linear Schroedinger equation. A topological connectome surface network can produce brain-like signals. For the Hodgkin-Huxley signals the details of the brain membrane played an essential role in modelling signals, while in this topological approach the detailed topological nature of a connectome surface with its surfaces’ charges and spin-half particles plays a similar essential role. With this general overview of the process of signal generation in place we now proceed to define a Riemann surface and its associated Riemann theta function and then provide the technical details required.

## 3 Technical details

We proved that our surface network exactly captures the topological features of any brain connectome as a surface and stated that it can be represented by a mathematical Riemann surface with spin structure an unknown large genus. We now define a Riemann surface.

### 3.1 Riemann surface variables

A Riemann surface *Σ*_*g*_ of unknown genus *g* is a geometric representation a algebraic polynomial equation *P*_*g*_ (*y,z*) = 0 of two complex variables (*y,z*). It can be represented as a collection of *g* linked doughnut surfaces as we stated before. The smoothness of the surface follows from the fact that the surface is constructed from a polynomial equation [29].

A genus *g* Riemann surface has two properties: it has topological connectivity and it has smoothness. The topological connectivity property can be captured by loop coordinates round the surface of the *g* connected doughnuts. There is a set of 2*g* of natural topological closed loop coordinates (*a*_*i*_, *b*_*i*_) that are used to represent the system’s connectivity. The *a*_*i*_ loops go round the tubes while the *b*_*i*_ loops go round the central hole of the *i*^*th*^ doughnut as shown (Fig. 1). These loops reflect the multiple periodic nature of *Σ*_*g*_. The sets of loops are said to be orthogonal. The smoothness of *Σ*_*g*_ is captured by *g* mathematical objects called one-forms *ω*_*i*_(*z*)*dz, i* = 1, 2, …*g*. The existence of these *g* one-forms was proved by Riemann [32]. A one-form is an object that can be integrated. Its smoothness is captured by the smoothness of the coefficients *ω*_*i*_(*z*), where *z* is a point on *Σ*_*g*_.

Thus *Σ*_*g*_, as a mathematical Riemann surface, can be described by the choice of its 2*g* topological coordinates (*a*_*i*_, *b*_*i*_) that describe closed loops and its smoothness is described by a specific set of *g* smooth one-forms.

### 3.2 The Riemann theta function

We next define the Riemann theta function Θ_*g*_ [28]. This function will use to study responses of the surface network due to input pinch deforming signals. We first define its variables and then define the function. The variables and parameters of Θ_*g*_ are constructed from those of *Σ*_*g*_ and its spin structure. We start with the parameters. An important set of parameters of Θ_*g*_ are those present in a symmetric matrix Ω_*ij*_, with positive imaginary values, called the period matrix Ω_*ij*_ which is defined by integrating the one forms *ω*_*j*_(*z*)*dz* over the closed loop *b*_*i*_. We have 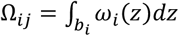, *i, j* = 1, 2, …*g*. It is a set of 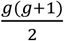 complex numbers.

The second set of parameters of Θ_*g*_ are a set of positive or negative integers that can range between (±∞). The integers come from integrating one forms *ω*_*j*_(*z*)*dz* over closed the *a*_*i*_ loops. These integrals 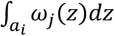 are normalized to be one when *i* = *j* and are zero otherwise. Thus, they produce positive or negative integer values as they “wind” round the *ai* loops of the Riemann surface. The winding numbers *n*_*i*_ count how many times the one-form integral loop *a*_*i*_ is traversed in the clockwise or anticlockwise direction. We next turn to the variables of the theta function.

Θ_*g*_ has *g* complex variables *z*_*i*_ that are generated from the one complex variable *z* used to define a point on *Σ*_*g*_. They are given by 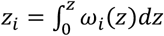. In words *z*_*i*_are generated by integrating the *i*^*th*^ one form between a fixed point 0 on *Σ*_*g*_ to a variable point *z* on its surface (Fig. 2A).

Finally, Θ_*g*_ has, as stated before, 2*g* discrete variables called its characteristics, (*α*_*i*_, *β*_*i*_). *i* = 1, 2, …*g*, that describe the spin structure of *Σ*_*g*_, in a precise way. Topologically distinct spin structures of *Σ*_*g*_ correspond to different choices of characteristic values. We will make this link clear shortly. Each characteristic can take one of two values, namely zero or half.

We can now define the spin topology integer *W* of in terms of the characteristics of Θ_*g*_, we have, 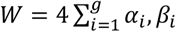. Thus, the spin structure topology number is the sum of the product of characteristics multiplied by four. The geometric meaning of these parameters will become clear once we write down the expression for Θ_*g*_.

Then its link to the spin structure of *Σ*_*g*_ will become clear. We now define the Riemann theta function 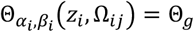

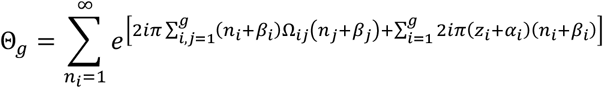

Notice the characteristics are related to the loops of *Σ*_*g*_ traversing a loop. It can be given a geometric interpretation produces. A traversal round a loop adds 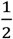 to if the spin is aligned to the surface and is zero otherwise, while the integer numbers *n*_*i*_ represent windings of the one-forms *ω*_*i*_(*z*)*dz* round *a*_*i*_ loops. The emergence of 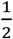 when an *a*_*i*_ loop is completed means the system has spin-half surface particles as a spin-half particle is a quantum object which changes sign under 2*π* rotation. This geometric result links characteristics of the theta function to spin structures on *Σ*_*g*_.

Thus we can represent the topological features of *Σ*_*g*_ by two topological numbers (*g,W*). The first *g* represents the connectivity of *Σ*_*g*_ and the second *W* describes its spin structure.

There is another way to think about the *g* variable points of Θ_*g*_ . The formula 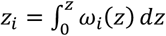 is now interpreted as a map between *Σ*_*g*_, a space of one complex variable with smoothness and periodicity properties, and a space of *g* variables with periodicity properties. The periodicity properties of this space come from the periodicity properties of these variables induced by the (*a*_*i*_, *b*_*i*_) loops of *Σ*_*g*_. This space is called the Jacobian *J*(Σ_*g*_) of Σ_*g*_. It provides an alternative description of *Σ*_*g*_ [28]

There is a map called the Jacobi Inversion map from *J*(Σ_*g*_ ) to *Σ*_*g*_ [28, 33]. The Riemann theta function is then defined as a function on *J*(Σ_*g*_). In our case we want the Θ_*g*_ to describe excitations of *Σ*_*g*_ so that its variables explicitly describe events on *Σ*_*g*_. This happens under suitable local topology changing surface deformations, input signals for signal generation.

We now have a mathematical description of *Σ*_*g*_ and of the function Θ_*g*_ that we will use to describe responses of the Riemann surface to pinch deformation input signals.

We next properly define the universal local topology changing input signals of Mumford, explain why theta functions have to satisfy the Fay identity and state the dynamical law in its final operational form that leads to Previato’s multi-soliton solutions [34]. This solution is then numerically evaluated for the case of two spikes and compared to an observed two action potential doublet recorded from a thalamic neuron. These results will complete the signal generating part of the paper. After that we move on to discuss memory formation, memory stability and then carry out three separate calculations that support the identification of soliton pulses as action potentials and shed light on the internal consistency of the method. At the end, in the discussion section, we will show that the network requires membrane gates to function properly and that the approach has points of overlap with existing brain modelling methods.

### 3.3 Input signals: Local pinch deformations

A pinch deformation locally reduces the circumference of a tube of the network to zero and as shown (Fig. 2), change its topology. Such a deformation can be described in mathematical terms by using of the *g* one-forms of *Σ*_*g*_. However, we can intuitively understand the effect of a pinch deformation on the *g* variables *z*_*i*_ of Θ_*g*_. Recall 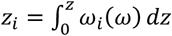. In the pinch limit the points (0,*z*) approach each other, where *z* is a point on *Σ*_*g*_ and the arc becomes a circle, whose radius approaches zero. Thus all *z*_*i*_ points converge to the one point *z* and this point itself approaches the centre of the tube that is pinched and we have *z*_*i*_ → *V*_*i*_*x* + *W*_*i*_*t* + *R*_*i*_ where (*x,t*) vary while (*V*_*i*_, *W*_*i*_, *R*_*i*_) represent the deformation. They are determined from the mathematical structure of *Σ*_*g*_. Let us explain how this is done. Consider the expressions 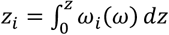. This is an integral of *g* one-forms between a fixed point 0 and a variable point *z* of *Σ*_*g*_. We now choose the two points (0,*z*) that are on a specific loop *a*_*i*_. A local pinch deformation at the point *z* corresponds to taking the limit *z* → 0 and at the same time reducing the radius R of this loop to zero. In this limit, 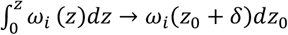, where *z*_0_ is the centre of the tube (Fig. 2A).

Expanding the one form coefficient *ω*_*i*_(*z*) near *z*_*0*_ then introduces parameters that represent a specific input local pinch deformation signal. The unknown parameter values define the self-generated information code that describes the pinch deformation. The precise mathematical details of this description are given in the Appendix.

### 3.4 The origin of the Fay trisecant identity

We now explain why a Riemann theta function has to satisfy the non-linear algebraic Fay trisecant identity if it is to be associated with a Riemann surface. It was proved [32] that a Riemann surface, *Σ*_*g*_ is uniquely specified by (3*g*−3) complex parameters, called its moduli parameters. These numbers essentially fix the shape of *Σ*_*g*_ but not its surface area or its volume in three dimensions [32]. They are thus topological in nature. Areas and volumes of geometric structures change are not topological variables as they change under continuous deformations.

We next note that the Riemann Theta function Θ_*g*_ is fixed by its period matrix Ω_*ij*_ which is a symmetric *g* × *g* matrix with 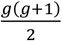 independent complex parameters. For *g* > 2 the number of parameters of Ω_*ij*_is greater than (3*g*−3). Thus, for *g* > 2 a Theta function with an arbitrary period matrix cannot represent a Riemann surface as it has more parameters than required. This means that if the Theta function is to be associated with a Riemann surface its period matrix parameters cannot be freely chosen: they must satisfy constraints. A remarkable result [28] was that if the Riemann Theta function satisfied non-linear Fay trisecant identity [35, 36] then it would represent a Riemann surface.

The problem of constraints can thus be solved in a practical way. We are now ready to state the dynamical law of the network in its final operational form.

### 3.5 Dynamical law of network: Final Form

> *The dynamical law for the network requires that during input pinch deformations the Riemann theta function must continue to satisfy the pinch deformed Fay identity and the input pinch deformations signals must respect the Riemann surface structure*.

We already pointed out that pinch deformation signals respect the Riemann surface structure as the deformation parameters are determined from the one-forms of the Riemann surface. Thus, requiring the theta function is satisfy the Fay identity is enough. The fundamental structural property of the network is preserved during signal production.

## 4 Results

We have given a conceptual picture of signal production. The dynamical law and input pinch deformations, with the help of the deformed Fay identity require output responses to be soliton solutions of non-linear differential equations: local pinch deformation change the topology of a signal producing subunit to a that of a sphere and they deform the Fay identity to become a nonlinear differential equation. But there is one remaining technical twist. We require that the Riemann surface of the network must represent a hyperelliptic equation. We next explain why.

### 4.1 Importance of the hyperelliptic equation

We would like pinch deformations to generate a wide variety of observed brain like signals using a previous discovery [28] but using his approach on a general Riemann surface does not lead to the variety of brain-like signals we want. But we do know that solutions of the one-dimensional nonlinear Schroedinger equation do have the desired variety of brain-like excitations we want. The non-linear Schroedinger differential equation can emerge as the pinch deformation equation of the Fay identity [30], only if the network surface is chosen to be the special Riemann surface that represents the hyperelliptic equation^1^, defined by the equation

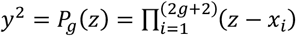, where *z* is complex and the roots *x*_*i*_ of *P*_*g*_(*z*) are either all real or if there is a complex root its complex conjugate must also be a root, then this leads to excitations that are Θ_*g*_ function solutions of the non-linear Schroedinger differential equation. The technical reason for this result is that the hyperelliptic equation produces a Riemann surface of genus *g* with 2*g*, not 3*g*−3 moduli parameters.

### 4.2 Signal generation

Let us summarise what we have established. We constructed a surface network with surface spin-half particles. We proved that the surface of the network could be chosen to have the topological connectivity of any brain connectome, could be represented by a mathematical Riemann surface with a spin structure and, if the Riemann surface was chosen to represent the hyperelliptic equation, then by local, topology changing, pinch deformation, signals, a wide variety of brain-like signals that were solutions of the non-linear Schroedinger equation could be generated. These result follow from the dynamical law of the network. We have thus established the result we were looking for. Brain-like signals are solutions of the non-linear Schroedinger equation, have pinch deformed variables and carry the topology numbers (*k,w*) of a signal producing subunit Σ_*k*_ and spin structure *w*. The process of signal generation reduces Σ_*k*_ to have the topology of a sphere (Fig. 2B and 2C), which has topology numbers (*k* = 0, *w* = 0) We now provide the final set of technical details regarding the solutions. We show that in the pinch limit all the *g* variables of Θ_*g*_ collapse to two one real variables (*x,t*) associated with one point *z* of Σ_*g*_ where (*x,t*) are multiplied by pinch deformation variables, namely, *z*_*i*_ → *V*_*i*_*x* + *W*_*i*_*t* + *R*_*i*_ where the parameters (*V*_*i*_, *W*_*i*_, *R*_*i*_) are fixed by the pinch deformation. The response to a pinch deformation are one-dimensional excitations *ψ*_*j*_(*x, t*) in Σ_*g*_ that are ratios of Θ_*g*_ function with the pinch deformed variables described.

Let us write down the non-linear Schroedinger equation when a subunit of genus *k* is pinch deformed.

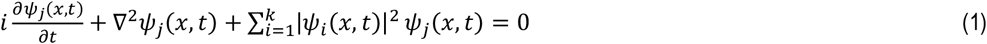

Here each *ψ*_*j*_(*x, t*) is a soliton pulse. It is a ratio of Θ function. The solution, in the soliton limit, has the structure 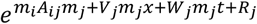 where *V*_*i*_, *W*_*i*_, and *R*_*i*_ depend on pinch deformation parameters. These parameters could then become a sort of the self-generated unknown information code of the network. The next step is to obtain the soliton limit of the solutions. We will use the soliton solutions previously found [34] to fit an observed doublet of action potentials recorded from a thalamic neuron. We will then comment on the fits and the numerical results obtained.

### 4.3 The soliton limit

Soliton solutions emerge in the limit when the genus *g* signal producing surface, collapses to genus zero. The technical way of doing this has been discussed [31] with numerical evaluation of localised transient excitations. We will not use these solutions as they have extra oscillation parts, but a different option [34].

#### 4.3.1 Dark soliton solution details [34]

We next write down the expression for the multi-soliton solution that we will use to fit observed two spike data for thalamic neurons. These solitons travel finite distances. The number of spikes *N* in the solution is *g* the genus of the signal producing subunit and the solution is multiplied by 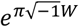 where *W* is the spin topology number of the subunit. It thus carries the two topological numbers of the subunit.

The multi soliton solution is given by the expression,

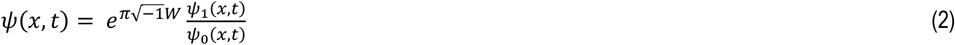

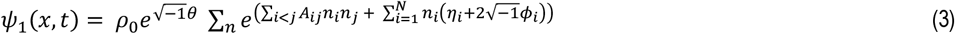

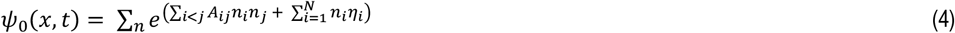

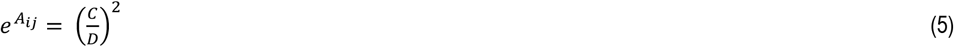

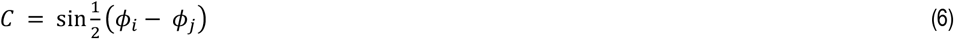

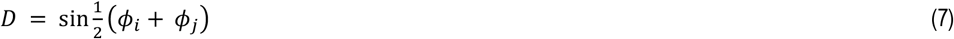

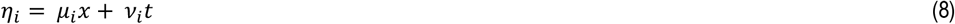

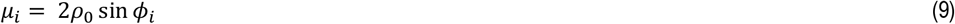

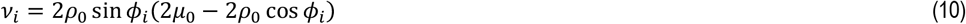

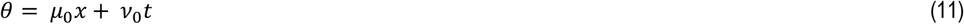

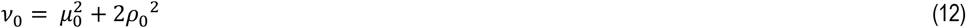

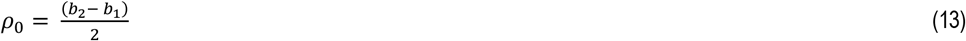

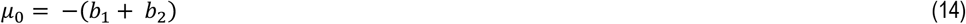

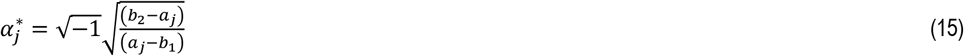

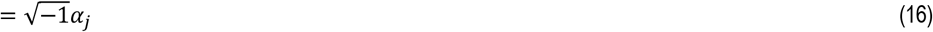

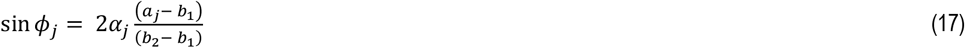

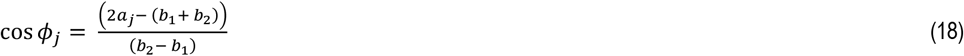

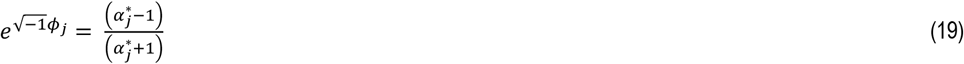

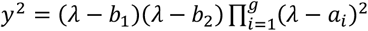 represents the pinched equation. The root entries are all real and are ordered as, *b*_1_< *a*_1_ < *a*_*2*_ ⋯ < *a*_*g*_ < *b*_*2*_

*N* = *g α*_*0*_ = −1.

### 4.2 Numerical results

We show that it is possible to fit the basic features of the observed thalamic neuron’s two action potential firing response to excitatory synaptic input (Fig. 3), such as their heights and separation distance between them by using the solution for *g* = 2. Each pulse has a different profile as predicted by the theory. The fits depend on the values of the four parameters (*b*_*l*_, *a*_*l*_, *a*_*2*_, *b*_*2*_) that define the two-soliton solution and require a choice for the variable *x*. A better fit is possible by adjusting the parameter values with greater care. But our point is simply to demonstrate that the essential properties of action potentials can be captured by soliton solutions.

**Figure 3.**
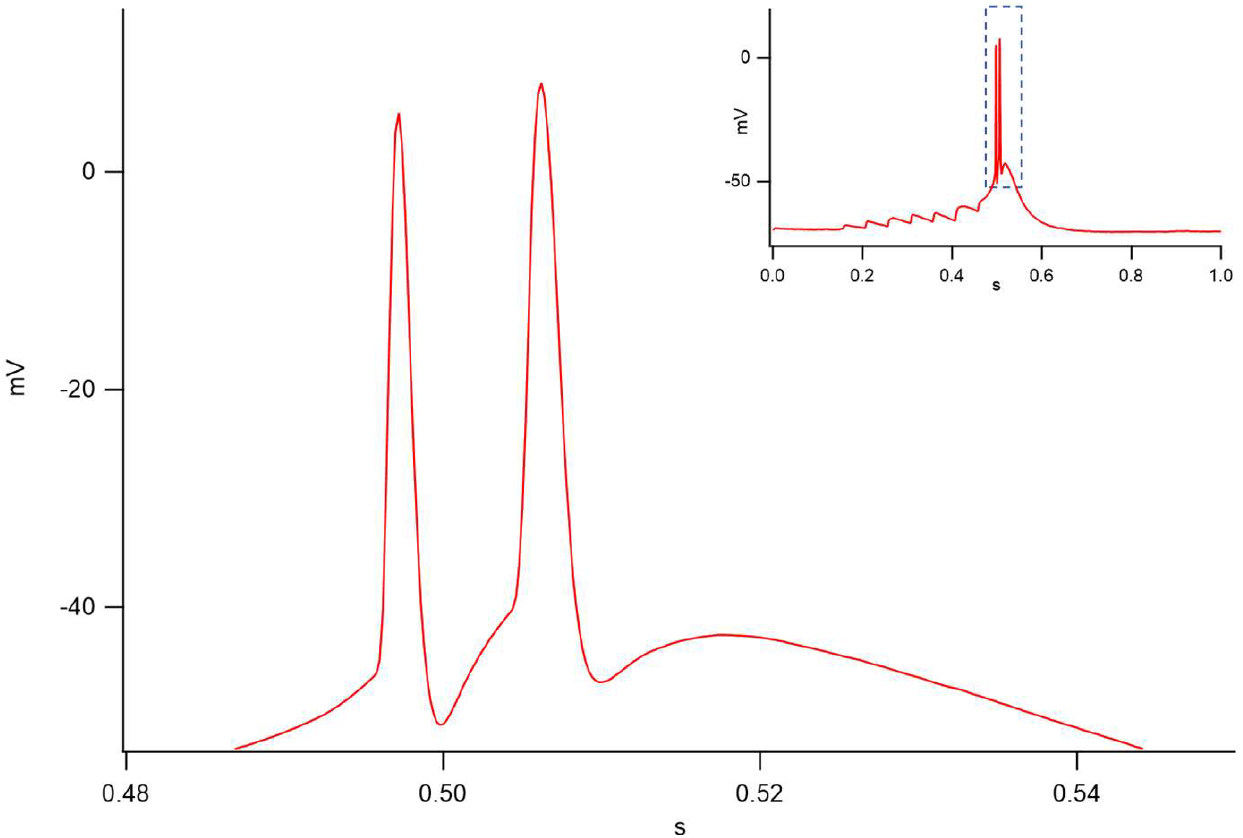
Detail of a whole cell current clamp recording (full; top right) of two elicited action potentials, from a murine thalamic (Po) neuron (IgorPro v9, Wavemetrics). Triggered as a response to presynaptic stimulation of corticothalamic axonal release, controlled by optogenetics. The protocol consisted in 8 consecutive 3ms light pulses (475nm) at 20Hz (details in Appendix).

**Figure 4.**
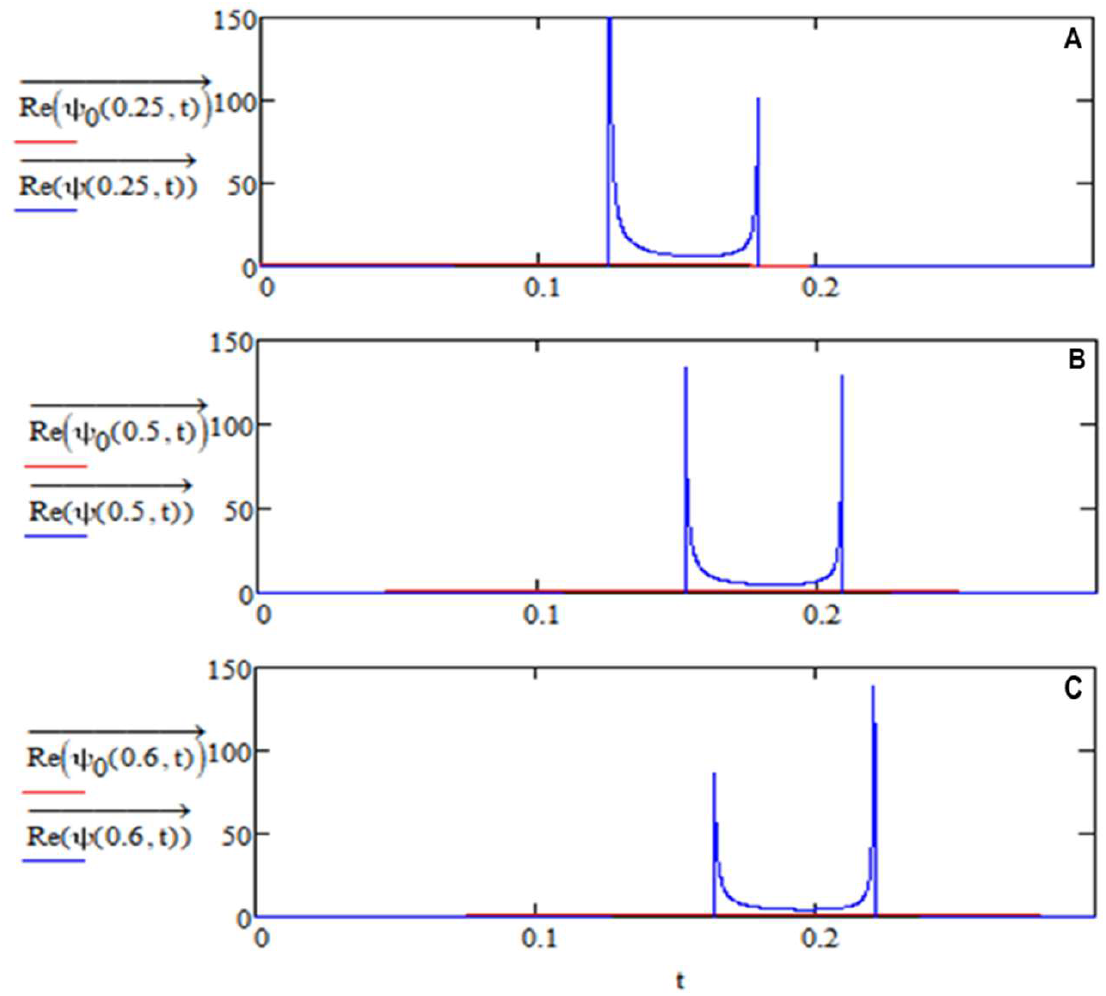
Three solutions for g = 2 with different spike amplitudes. We fit the first solution to the data by choosing time and voltage scale parameters (Mathcad 150, MathSoft).

**Figure 5.**
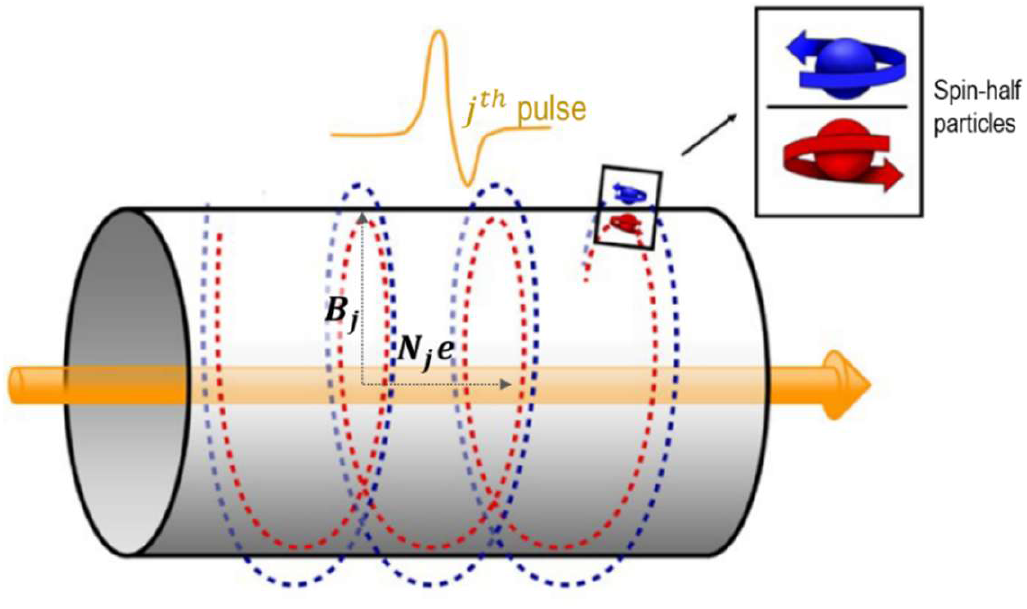
Helical structure produced by magnetic field *B*_*j*_ on the spin-half particles present on both sides of the tube surface by the advancing charge *N*_*j*_*e* carried by the *j*^*th*^ pulse.

**Table 1.** Parameters of soliton solutions with comments.

| SYMBOL | FUNCTIONS | VALUE | COMMENT |
| --- | --- | --- | --- |
| $b_1$ | Parameter | Input signal | Smallest real number |
| $b_2$ | Parameter | Output signal | Largest real number |
| $a_j$ | Parameter | Pinch Locations | Ordered real numbers |
| $\alpha_i$ | Function of parameters | Fixes Period Matrix | Imaginary numbers |
| $\phi_j$ | Function of $\alpha_j$ | Fixes Period Matrix | Real numbers |
| $N$ | Parameter | Number of Solitons | Equal to genus |

**Table 2.** Soliton parameters. Note: Previato’s solutions describes multiple pulses in one expression in which the rate at which pulses appear and their separations are all fixed by the input deformation parameters.

| **g** | **x** | **($b_1, a_1, a_2, b_2$)** | **Figure** |
| --- | --- | --- | --- |
| 2 | 0.25 | 0.3, 0.08, 1.05, 2.3 | 4.A |
| 2 | 0.5 | 0.3, 0.08, 1.05, 2.3 | 4.B |
| 2 | 0.6 | 0.3, 0.08, 1.05, 2.3 | 4.C |

A table summarising the parameters and functions for a *g* multi-soliton solution. All the other parameters that appear in the solution, such as, (*η*_*i*_, *ρ*_0_, *μ*_0_, *μ*_i_, *ν*_i_) are functions of the parameters listed in the table.

Their dependence on these parameters are given in the expression for the solutions. The ordering of the real parameters was also stated earlier. Namely (*b*_l_ < *a*_l_ < *a*_2_ < ⋯ < *a*_g_ < *b*_2_). For the two spike case *g* = 2 we thus have (*b*_l_ < *a*_l_ < *a*_2_ < *b*_2_) *b*1 and for a *g* = 4 we would have (*b*_l_ < *a*_l_ < *a*_2_ < *a*_3_ < *a*_4_ < *b*_2_). The two spike (*g* = 2) solution parameters used to numerically determine them are displayed in the table and their numerical plots are shown. The range of possible excitations is illustrated by three numerical examples in which the relative amplitudes of the two spikes are different.

For *g* = 2 the full expression of Eq.2 used is as follows:

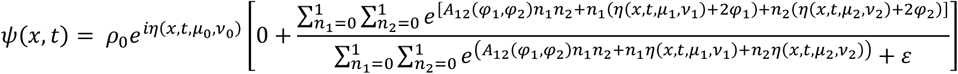

### 4.4 Comment on the solutions

As stated, and confirmed by our numerical results, a wide variety of brain-like signals can be generated by pinch deformation input signals, they include local transient excitations [37] as well as different propagating multi-soliton solutions [34]. These solutions are dimensionless, but as we now show, it is possible to choose units of length, time and voltage so that the solutions *ψ*(*x, t*), which represent action potentials, are in volts.

To see we first rewrite the non-linear Schroedinger equation by including two parameters (*D,h*)

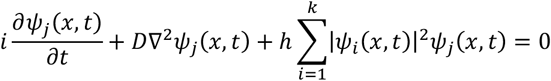

Then we introduce dimensions of length *x*_0_ of time *t*_0_ and voltage *V*_0_ and write our dimensionless variables (*x,t*) and the dimensionless function *ψ* as: 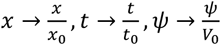. These dimensions factors modify (*D,h*) to 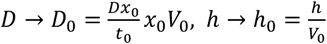. If we now set (*D*_0_ = 1, *h*_0_ = 1), we get back our equation but now with solutions *ψ* that are voltages. The dimensional value choices for (*D*_0_, *h*_0_) define the equation.

The theoretical method of generating soliton signals by local pinch deformation driven by mathematics [28] is supported by biology. It is observed that pinch deformations of an axon tube produce action potentials [38].

We point out that the method of signal generation described for a signal producing subunit *Σ*_*k*_ only works if the characteristics of the corresponding Riemann theta function Θ_*k*_ are not zero and *e*^*iπW*^ = −1, where 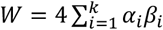. is the spin topology number and *k* the genus of the signal producing subunit. Because if this condition is not satisfied the Fay identity does not exist hence the method of signal generation described does not work. Non-zero characteristics represent spin structures. Thus, surface spin-half particles are essential for generating signals. We next show how propagating signals produced store memories in the pathways of the network that they traverse as claimed.

### 4.5 How are memories are stored in the network?

Propagating soliton pulses carry information of their creation with them. This follows from the fact that the variables of the Riemann theta function are derived from the variables of its associated Riemann surface. Thus, a pinch deformation input signal of the surface of the Riemann surface induces deformations of the theta function variables of the theta function output signals. We now show that this information can be transferred by the moving voltage pulse signals to align spin-half surface particle present in the pathways traversed by the solitons to create a memory structure. The memory structure is naturally created by the laws of electromagnetism. Moving soliton pulses carry charges and produce helical magnetic fields, as they move [21] that act on surface spin-half particles and align them to create a helical linked spin-half memory structure. To study this process quantum theory has to be used as spin-half systems obey laws of quantum theory. The aligning energy is the Hamiltonian of the system and is given by the operator expression 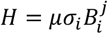. where 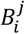 is the magnetic field generated by the *j*^*th*^ pulse, and *μσ*_*i*_, *i* = 1, 2, 3 is the spin magnet with *σ*_*i*_ the quantum description of spin in terms of matrices *σ*_*i*_ known as Pauli spin matrices [39, 40]. The Hamiltonian operator *H* defined determines the time evolution of the spin [39] and produces a helical spin memory structure. What is not clear is if the structure created is stable under thermal disruptions. This requires analysis. It is shown that provided certain conditions on the number of pulses present in a signal and the charges they carry are met the structures formed are stable. They are memories. If the conditions are not satisfied the memory formed is not stable. It represents a short-term memory. We will prove that each memory structure has a specific excitation frequency and suggest that this frequency is used to retrieve it.

Let us sketch the mathematical steps (more details are in the Appendix). The alignment of surface spin due to the magnetic field is a quantum process^2^ . If the initial spin-half particle state vector is *S*(0) at time *t* = 0, its time evolution to a state *S*(*t*) is given by the equation

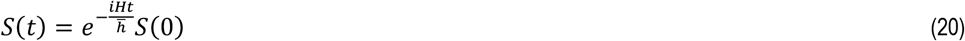

where the Hamiltonian *H* is defined as 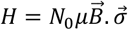 represents the interaction of spins, described by Pauli matrices *σ*^_*i*_^, *i* = 1, 2, 3 with an external magnetic field 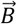. The magnitude of the magnetic field is 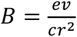, with *c* the velocity of light, *v* the speed of the soliton pulse, *N*_0_*e* the charge of the pulse, 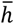, Planck’s constant and *r* is the radius of axon tube.

A quantum calculation, (see Appendix) shows that the helical spin-magnet structure created by the transient pulse magnetic fields of moving solitons, is given by,

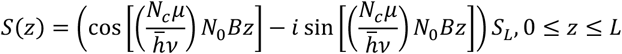

where *N*_0_ is the number of electrons in the pulse, *S*_*L*_ is *S*_*i*_(0), the initial spin orientations values at *z* = 0, *L* is the length of the memory structure and *N*_*c*_ proton spins located on a circle with centre *z*. These spins are aligned at the same time. Thus, the magnetic field acts on all *N*_*c*_ spins at the same time. The magnetic field *B*_*j*_ produced by an action potential pulse has been measured [42] and it has been shown that it can be used to accurately reconstruct the action potential. The spin aligned structure contains the magnetic field details it thus has all the information carried by the soliton voltage pulses. It is therefore a faithful recording of the information carried by the soliton pulses. If it is stable, it is a non-transient true memory structure.

In our discussion we ignored the detailed nature of the magnetic field generated by the *j*^*th*^ soliton pulse and simply represent its magnitude by the expression 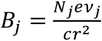, where *r* is the radius of the axon, *v j* the speed of the soliton pulse, *e* the electric charge, *c* the speed of light in the medium and *N*_*j*_ the number of units of charge carried by the soliton pulse. The direction of the magnetic field is tangential to the axon surface circle of radius *r*. These simplifications allowed us to write down explicit expressions which illustrate the memory structure and its formation.

Memory structure formation is thus a natural process. Its existence and stability, however, are matters that need investigation. After formation the stability of the structure requires that the total binding energy of *N*_*L*_ units of binding energies between neighbouring spins on the helical memory structure can withstand thermal disruption energy *kT*. Thus, we require binding energy expression *E* > *kT*, where 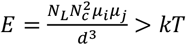 where *d* is the distance between the two neighbouring spins, *k* is the Boltzmann constant, *T* the temperature in degrees Kelvin, *N*_*c*_ is number of spins on neighbouring circles of the helical structure, and *N*_*L*_ is the number of such paired spins in the full memory structure, which behaves as a single unit under thermal disruptions. If this condition is satisfied, a stable long-term memory is created, otherwise the memory is short term. The issues of the stability of memories are discussed in greater detail elsewhere [41]. Putting in numbers we have 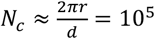, *d* ≈ 10^−8^*cm, N*_*L*_ ≈ 10^8^ per cm, *k* ≈ 10^−16^*ergs*/*deg, T* ≈ 100°*C*. Thus, a memory structure of length greater that 10^−3^cm will be stable. We now point out that the helical memory structure does not have net macroscopic magnetic properties, because two neighbouring spin-half particles on a helical structure have opposite orientation under 2*π* radian rotations. This is a special quantum property of spin-half particles.

#### 4.5.1 Memory excitation frequency estimate

Each memory is found to have a characteristic excitation frequency of the structure *ω*_*j*_ that can be estimated by the formula 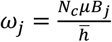, It depends on 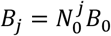, where 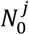 is the charge carried by the *j*^*th*^ pulse and 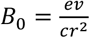 is the magnetic field created on the axon surface by a unit charge moving with speed *v*. The memory excitation frequency is signal specific. The details are in the Appendix. We find that 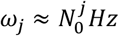.

#### 4.5.2 Memory recall

The storing of memories on the surface of axon tubes by aligning spin-half particles, is as we saw a natural physics driven process. It depends on two facts: soliton pulses carry charge and thus generate magnetic fields and these fields naturally align surface spins as a helical structure, which are the memory substrate. Each memory has a specific excitation frequency in the range 1−20Hz that we suggest can be used to retrieve it by a resonance excitation method. Our memory excitation frequency estimate assumes the memory structure formed is due to spin half protons. The reasons for this choice are discussed elsewhere [41]. The structure is consistent with the picture of memory engrams [16].

## 5 Solitons as action potentials

We identify observed action potentials with soliton pulses. The identification is supported by a number of checks. The first check has already been made. This was to fit observed action potential spikes to soliton solutions [34]. The next set of checks examine if the network can accommodate well known properties of action potentials such as their resting potential value. We now show that this is possible.

### 5.1 Estimate of the value of the resting potential

The standard method for estimating the resting potential value in biology is by using the generalized Nernst equation for chemical processes based on equilibrium thermodynamics [43]. The estimate is an approximation as the axon membranes are never in equilibrium but are constantly involved in the transmission of signals. In this approach the flow of surface sodium ions into the membrane tube and the outflow of potassium ions due to the opening and closing of membrane gates is regarded as an equilibrium thermodynamic process and method of chemical potentials of thermodynamics is used to determine their equilibrium potential values [3, 44]. Our approach is different. We will calculate the value of the resting potential by considering two strips of the axon membrane surface separated by the diameter *r*, using the electrical properties of the membrane with the help of the quantum electrodynamic calculations [18]. The calculation shows that there is a dynamic attractive Casimir-Polder potential *V*_*CP*_(*r*), between these two charged dipoles layers separated by a distance *r*. The idea is that this attractive potential is balanced by the elastic properties of the tube and the fluid inside the tube that fix the value of *r* and thus the dynamic equilibrium value of the resting potential is the value of *V*_*CP*_(*r*), with *r* set equal to the observed axon radius. The value of *r*, as an experimental input, that gives our result. We have [18],

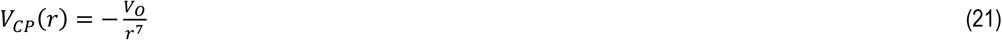

which is an attractive potential energy between dynamic neutral molecules. The factor *V*_O_ has an explicit form that we do not use. The resting potential value of the axon membrane tube determined does not require knowledge of the details of the fluids inside the tube. An Important feature of the Casimir Polder potential is that the result of zero-point energy. It does not depend on the electric charge *e* but on the combination 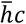, the Plank constant and the speed of light.

The problem is to estimate the value of *V*_*O*_, for a fixed experimental input value of *r* using the membranes dynamic dipole nature and thus find the value of *V*_*CP*_. We estimate a value of ≈ 70mV simply using dimensional reasoning.

Let *d* be the membrane thickness, *r* the diameter of the tube, *e* the charge of an electron, *r*_*b*_ the hydrogen atom Bohr radius, and *N* the effective number of molecules in the segment of the membrane chosen. These variables, other than *N*, are taken to have the following values [45]:

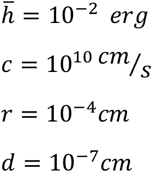

We next write *V*_*O*_ as

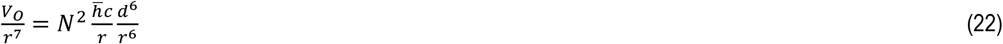

*N* is determined by setting *V*_*CP*_(*r*) = *V*_*E*_, where *V*_*E*_ is the resting potential.

Thus, we have

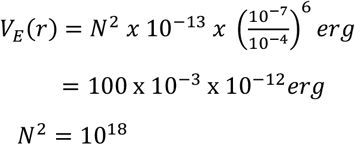

For the idea to work, the value of *N* should be *N* ≈ 10^lO^, where *N* is the number of dipoles in a segment area of width *w* and length *L*. The observed number of dipoles per square per square on a membrane is known [46] and can be ≈ 10^14^ per cm^2^. Using this number that the area segment that contain *N* dipoles has *w* ≈ 10^-^ cm and *L* ≈ 10^-l^cm

### 5.2 Observable consequences of network

We next calculate two observable consequences of the approach that suggest that membrane ion channels are required for soliton pulse propagations. The first consequence is the that membrane distortion waves are expected to accompany soliton pulses as they propagate along axons. The next is a theoretical estimate for the separation distance between the nodes of Ranvier. The nodes of Ranvier are gaps in the myelin sheath cover of axons that are observed [20].

We now derive a formula relating a soliton pulse action potential *A*(*x,t*), and its accompanying surface distortion wave Δ(*x,t*).

### 5.3 A Formula relating Δ(*x,t*) **to** *A*(*x,t*)

Our formula follows from two inputs: a mathematical link between geometric distortions of our surface tube and voltage changes, and from the use of low Reynold numbers physics for transverse motions of the tube. The first input has already been used. It is the Casimir Polder potential. The second input can be easily justified. The Reynolds number *R* of a fluid is given by 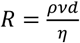, where *v* is the transverse fluid velocity, *d* is the diameter of the tube, *ρ* the fluid density and *η* is the dynamic viscosity coefficient of the fluid. In the transverse direction *d* ≈ 10^−4^cm and *v* is small so that *R* is small. In a low Reynolds number environment, there are no time delays and no inertial effects [19] so that a voltage changes directly translates to deformations in the transverse direction to the propagating voltage pulse Thus, the Reynolds number physics implies that the membrane distortion should be proportional to the soliton pulse action potential. We have,

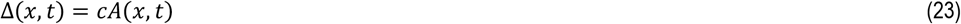

We have to fix the proportionality factor *c*. We do this by using the Casimir Polder to relate a membrane potential change Δ*V* to a membrane distortion. For the distortion *r* → *r*_*E*_ + Δ(*x, t*) where *r*_*E*_ is the equilibrium axon diameter, the corresponding Δ*V* is,

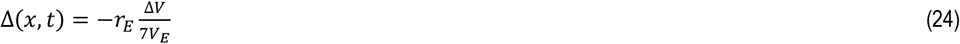

where *V*_*E*_ is the equilibrium membrane potential. We note that Δ*V* is zero when there is no distortion and the membrane is at its resting potential value of *V*_*E*_. We now chose *c* so that it agrees with the result for Δ*V* when *A*(*x, t*) = Δ*V* = |*V*_*T*_| − |*V*_*E*_| for some value of (*x, t*). This input fixes the value of *c*, and gives the formula for the propagating membrane distortion wave,

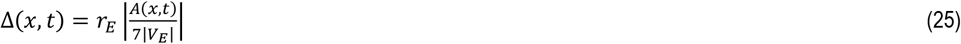

We next set (*x, t*) = *z* = *x* − *vt*. Then *D*(*x, t*) = *D*(*z*) and *A*(*x, t*) = *A*(*z*). For qualitative understanding we represent *A(z)* by the expression,

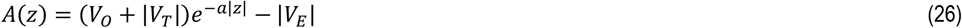

where *v* is the speed of the pulse, *V*_*M*_ its maximum is reached when *z = 0* and *-1 < z < +1, a* > 0. We have two conditions *A*(0) = *V*_0_ + *V*_*T*_ + |*V*_*E*_ | = *V*_*M*_ and (*V*_0_ + |*V* _*T*_ |)*e*^-a^ = |*V*_*E*_ |. This gives 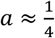. Using the diameter of the axon as a scale for *z*, in keeping with the Reynolds number assumption of modelling only transverse effects, gives a width of the action potential of ≈ *0*.*4µm*.

The theoretical formula thus, makes two predictions: the distortion wave amplitudes are proportional to the action potential and their speeds are those of the action potential. We can estimate the maximum distortion amplitude by setting *r*_*E*_ ≈ *10*^−*5*^*cm, V*_*T*_ ≈ −*65mV, V*_*E*_ ≈ −*80mV*, and the range −*65 < A(z) < +20mV* for the rat’s hippocampus axon radius [47]. The formula then predicts that the distortion wave will have a maximum value of *Δ*_*M*_ ≈ *30nm*. The threshold potential value *-65mV* is an estimate. The essential point of the analysis was to establish the proportionality of the distortion wave to the action potential using low Reynold numbers physics.

### 5.4 Distance between the nodes of Ranvier

It is known that axons are covered by insulating myelin sheaths, but there are nodal regions that are not covered. These are the nodes of Ranvier [20]. They are regarded as signal boosting regions. We will show that this is true and will explain the way the signals are boosted. Our suggested method of boosting signals makes use of different results established for the surface network.

We estimate the energy lost by soliton pulses by aligning surface proton spins to form memory structures. We then estimate the energy loss of a soliton, per cm of a multiple soliton pulse that can create a memory. It is required that this energy loss after traversing a distance *L* should be equal to the energy of an action potential pulse. This explains why signal boosting is required and the observed gaps between the nodes tells us the amount of energy loss allowed before the signal is boosted. It thus depends on the choice made by the brain to boost signals after a certain percentage of energy loss. Observationally energy loss of soliton pulses should show up as reduction of their amplitudes.

The observed length between the nodes is obtained if signal boosting happen after an energy loss of ≈ 1% .

### 5.5 Estimating the gaps between the nodes

We will estimate the spacing between nodes in the following way. We will calculate the energy loss of one soliton pulse per cm of travel due to the energy it spends aligning spins. The point to note is that even though solitons are non-dissipative excitations they can lose energy if they are involved in interactions. In this case the soliton pulses interact with surface spins. The interaction energy is through the magnetic field generated by the soliton and is given by *E = N*_*c*_*μB* where *μ* is the proton magnetron [41]. The multiplicate factor *N*_*c*_ represents the number of spins that are aligned at the same time by the helical field. It is given by the number of protons on a circle of radius *r* surrounding the axon tube. Thus 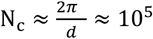 where *r* ≈ 10^−4^cm and *d* is the separation distance between protons, *d* ≈ 2.5×10^−8^cm.

To pin down the energy loss we need an expression for the magnitude of the helical magnetic field *B* generated by the moving soliton pulse of charge, *N*_0_*e*, where *e* is an electron charge, and speed *v*. It is given by *B = N*_*0*_*B*_*0*_ where 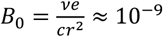 gauss, *r* is the radius of the axon and c the speed of light.

As we are interested in the energy loss of the soliton pulse per cm of propagation, we need an estimate for the number of protons present in this length. The magnetic field is helical, it acts on all spins present on the nerve surface above its trajectory. Thus, if *N* number of proton spins per square cm, then the number of surface protons present for unit length of the axon tube is *2πr × l × N* with *l* = 1cm. The energy loss per cm length of one soliton pulse, is thus given by,

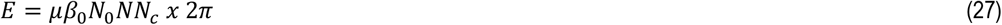

For length *L* the energy loss is then *E*_*L*_ = *LE*. We want to find the length *L* the soliton pulse needs to travel to lose all of its energy. This condition gives the equation *LE* = *N*_0_*eV*, where *N*_0_*eV* is the energy of the pulse, *V* its voltage and *N*_0_*e* it’s charge. This gives a formula for *L*,

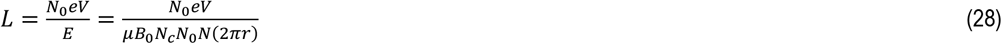

Let us put in numbers, *N eV* ≈ *N* 10^−13^ergs, *μ* ≈ 10^−23^, *N*_c_ ≈ 10^5^. We get 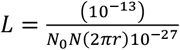,. We will find that *N*_0_*N*(2*πr*) ≈ 10^14^, so that we get *L* = 10cm. Note that the value of *N*_0_ plays no role in the calculation. We give the estimate details: 2*πrN* ≈ 10^13^, since the area swept out when the pulse travels one cm distance along the axon tube is 10^−3^ × 10^16^ = 10^13^. Thus, we find that a pulse loses all of its energy after ravelling a distance *L* = 10cm. For length ≈ 10^−1^cm the energy loss of the pulse is 1%. Making this choice gives us an estimate for the separation distance between the nodes of Ranvier of ≈ 10^−1^cm. This is roughly the observed value. The estimate is crude but it makes a prediction that after travelling a distance *L* ≈ 10cm a soliton pulse loses all of its energy^3^. The calculation makes use of a number of results of the network and suggests a universal method of boosting signals: it uses the idea that memory structures involve aligning spins which, by the laws of physics, drains energy from the soliton, it uses the idea that pinch deformations generate signals and it uses the result a membrane distortion wave amplitude is proportional to the amplitude of the soliton pulse that creates it.

We now make two comments. When we discussed the stability of memory structures, we found they could not be produced by a single pulse but require multiple pulses for their creation. We can extend the calculation for gap separation to consider the energy loss of multiple pulses but such a calculation will not change the estimate of the gap separation since for *N*_*s*_ pulses both sides of our equation, becomes *N*_*s*_*EL = N*_*s*_*N*_*0*_*eV* and the factor *N*_*s*_ has no effect on the value of *L*. The second comment is the estimated bound for *L* of 10cm after which a soliton pulse loses all of its energy means that signal boosting is required. For myelinated nerve fibres we explained how this boosting happens at the nodes of Ranvier. But there are unmyelinated nerve fibres. Signal in them also require boosting. Some have long lengths. For example, C-fibres, that carry the sensation of temperature and pain with them, are unmyelinated and have lengths greater than 100cm. The theory predicts that boosting is possible using the same ideas described for myelinated nerve fibres, if all unmyelinated nerve fibres have diameters that vary. Then the small diameter regions of the nerve fibre are pinch regions. This prediction turns out to be correct. The diameter of the C-fibres does indeed vary (0.2-1.5*μm*) [48], which represents a considerable variation. But the spacing between regions with large and small diameters is not uniform. However, the mechanism of boosting is the same as for myelinated nerve fibres. In a pinch region of the unmyelinated nerve fibres, the correct pinch deformation parameters required to boost a given pulse without distortion, are provided by the modulations of the membrane deformation waves. We expect this mechanism is operative in all unmyelinated nerve fibres. Thus, this prediction suggests that membrane deformation waves play a key role in signal propagation.

## 6 Discussion

We started by raising a few fundamental unresolved conceptual problems of theoretical neuroscience. The first was that action potentials signals cannot carry information regarding their creation. The other issues were a lack of theoretical understanding of how and where memories are stored and how they can be retrieved. As action potentials do not carry information in current models there is no theoretical way to link signals to memory creation in them.

We claimed that these problems can be solved using an unconventional method of signal generation using a special surface network with surface spin-half particles. The claim has been demonstrated. We have shown that a special surface network with the topological connectivity of any brain connectome exists, has a mathematical representation as a Riemann surface with spin structure and could generate a wide variety of brain-like signals in response to topology changing pinch deformations. Physical pinch deformations are known to generate action potentials [38]. We showed that the signals carry information of their creation with them encoded in the network’s own self-generated code of pinch deformation parameter values. We then showed that there was a natural mechanism for the signals to transfer this information along the pathways they traversed, by aligning surface spin-half particles, to create memory structures. The alignment of spins follows from the laws of electromagnetism. A mechanism for recalling memory by resonance excitation was also suggested based on the theoretical result derived, that each memory structure has a specific excitation frequency.

Therefor we have presented the structure for a memory forming network. One that could align to our current understanding of neurobiological circuits and provide new possible insights. Where all possible inputs (e.g.: sensory) are converted to a common self-generated code of deformation parameter values associated with the pinch deformation.

Signal generation in the network is a global topological process. It requires the topology of a signal generating subunit surface with topological numbers (*k,w*) to be reduced, during signal generation, to a surface with topology numbers (*k* = 0, *w* = 0) that defines a sphere. What happens to these topology numbers? The answer is the signals carry these numbers with them. The signals are thus topological objects. Consequently, signal generating results remain valid under continuous deformations of the network and the geometric size of the network does not play a role in signal generation. Thus, it is expected that similar brain-like excitations can be produced by networks with volumes that can vary over a wide range of values.

Both topological numbers carried by signals are important. The connectivity, genus, number *k* shows up as the number of soliton pulses produced by the subunit, while the number *w* as we show, in a separate work [41], generate EEG waveforms and fixes their frequency and amplitude values. Here we showed that the geometric pinch deformation information carried by propagating signals are stored in memory structures. The stability of the memory structure created by signals requires investigation. We showed that if a structure was formed it would have thermal stability provided the size of the memory structure had more than e 10^5^ spin pair links. A thorough discussion of memory structure sizes, memory structure formation times, and their stability are given in a separate paper [41], where it is also conjectured that the memory structure is composed of spin-half protons. Reasons for this conjecture are given^4^.

At present there is no direct evidence of such helical spin-half magnetic structures in the brain but such structures have been observed in condensed matter physics [49, 50]. Finding the structure is difficult because, as we showed, the spin aligned memory structure, due to special quantum properties of spin-half particles, has no net macroscopic magnetism.

The consistency of the network picture and its ability to reproduce observational results of the brain was checked in a number of ways. The first test was to show that an observed two spike action potential could be fitted to a theoretical two soliton solutions. The next it was to show that a dynamical calculation of the membrane resting voltage value could be carried out to give a value that was in agreement with observations. Then a calculation that predicted the appearance of observed membrane distortion waves that accompany soliton pulses that correctly predict what is observed and gave a formula relating membrane distortion wave amplitudes to the amplitudes of the soliton pulses that generate them. Finally, a calculation estimating the separation distance between the nodes of Ranvier was carried in which the results of the other calculations as well as the pinch deformation method of signal production were all used to suggest a way soliton pules could be boosted without distortions. This calculation brought together different theoretical strands of the approach and produced a result that agreed with observational data. It suggested that unless boosted a signal will lose all of its energy after travelling 10cm. The calculation thus predicts that unmyelinated nerve fibres must have diameter values that vary with regions of low diameter that are natural pinch regions where a signal can be boosted by pinch deformation generated by membrane deformation waves.

Although the approach is global and non-reductionist certain features of standard theories are present in it and they play an important role. The first feature is that soliton pulse propagation requires the presence of membrane gates. An initial pinch opens a membrane gate and allows ions to flow in. As the soliton pulses propagate it produces a membrane wave distortion. When the membrane distortion waves expand the membrane ions flow in, when they contract the membrane ions flow out, in this way the flow of ion maintain the internal fluid pressure inside the axon tube so that no dissipative energy loss due to pressure differences creating dissipative fluid movements. Thus, the ion flows play and ion channels play an essential role in the propagation of soliton pulses but these flows are not the driving force for the propagation of the pulses: the driving force comes from topology of the network and the pinch deformations.

The approach also has common features present in neural mass models [52, 53] and in computational neuroscience models [4, 54, 43]. Neural mass models consider assembly of similar neurons. This feature is also present in the surface network as only an assembly of linked neurons described by a Riemann surface can generate signals. But unlike the neural mass model the interactions between assemblies of neurons are not ad-hoc inputs but are naturally generated by the non-linear topological structure of the network. Current computational neuroscience models use the brain connectome represented as a linear network in order to understand how signals propagate in the brain. Here too there is a convergence of ideas as the surface network makes use of the exact topological connectivity of a connectome, but now represented as a surface, to derive its results.

Finally, we stress that a major difference between the surface network and all existing models is that spin-half particles play an essential role both for signal production as well as for creating memory structures.

To conclude we emphasise that the surface network offers a different non reductionist topological way of thinking about brain functions where the structure of the network determines its functions. Further study of the approach seems worthwhile [41].

## ACKNOWLEDGEMENTS

The authors would like to thank Mike Coey, Kumar Gupta, Mani Ramaswami and Arnab Das for encouragement, Paul Voorheis and Mike Coey for reading through an earlier version of the manuscript and making helpful editorial suggestions, Raul Ramos for his help with the figures, Tim O’Leary for useful comments on earlier versions of the manuscript, Christian Klein and Jörg Frauendiener for their kind assistance and for supplying codes used in their work on Riemann surfaces, and also to Mauro Ferreira for his guidance and support.

## Author contributions

Siddhartha Sen carried out the formal analysis and wrote the paper. Tomas Ryan and Maurizio Pezzoli edited the paper and provided neuroscience inputs, Maurizio Pezzoli provided the thalamic spike data, contributed to the writing of the paper and supplied figures, David Muldowney contributed to numerical results and Plamen Stemanov fitted the spike data to soliton solutions and also produced different numerical soliton pulse solutions.

## There is new data used in the paper

The new data is due to Maurizio Pezzoli and is presented in the Appendix.

## Ethics

No ethical issues are involved in this work. All procedures to the animal were under the supervision and authorization of DGAV-affaires veterinaires du canton Vaud, Switzerland license VD3802, and according to EPFL veterinary regulations.

## Use of AI

No AI tools were used.

## Methodology and resources

The methodology used is publicly available. Maurizio Pezzoli’s method of data extraction is given in an Appendix and his data is available from him. Other resources used are referenced published papers and data available in the public domain.

## A Appendix

### 7.1 Hyperelliptic equation

We briefly sketch how to construct the Riemann surface *Σ*_*g*_ associated with the hyperelliptic equation and state some important properties of this Riemann surface.

#### 7.1.1 How is Σ_*g*_ constructed from *P*_*g*_(*z*)?

*Σ*_*g*_ is defined by its *g* one-forms and its 2*g* loops (*a*_*i*_, *b*_*i*_). We sketch how these are constructed from the hyperelliptic equation *y*^*2*^ = *P*_g_ (*z*). The *g* one forms are defined by the equation 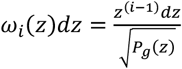 while the loop coordinates correspond to branch cuts between the zeros *x*_*i*_ of *P*_*g*_(*z*). We note that the zeros of *P*_*g*_(*z*) are real or appear as a complex conjugate pair so that y^2^ is always real.

We draw attention to two important properties of this *Σ*_*g*_: it can have (*g* + 1) subunits [55], giving it a modular structure and it has the maximum allowed number 2*g*, spin structures [56]. As we will relate spin structures to memory so that this is an important result as it places a bound on the number of memories that can be stored.

#### 7.1.2 Fay trisecant identity

For the sake of completeness, we briefly provide some mathematical details regarding the structure of the Fay identity in the pinch deformed limit [31]. The soliton solutions are found by using this identity. In the logic chart (Fig. 6), the intricate links between the Riemann theta function and Riemann surfaces is shown. In it the Fay trisecant identity is also displayed using a full notation [36].

**Figure 6.**
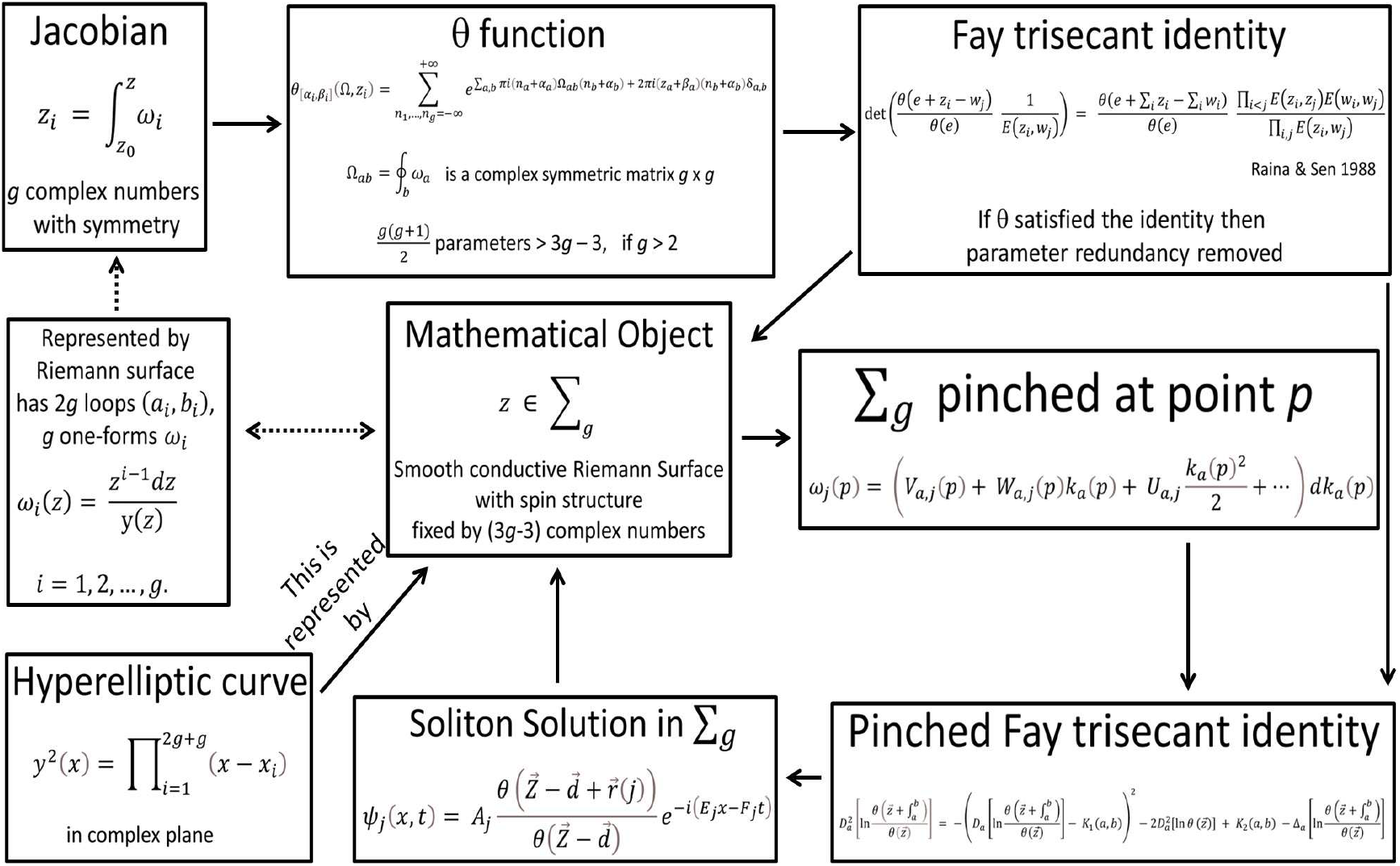
Mathematical logical chart of the conceptual interactions that conform the presented framework, the interaction of them through the dynamical law that allows us to solve it, generating soliton solutions.

We first define a number of functions and then use them to write down pinch deformed form of the Fay identity. Let (*a, b*) be distinct points on *Σ*_*g*_. We fix local parameters *k*_*a*_, *k*_*b*_in a neighbourhood of these points. Let *δ* = (*α*_*i*_, *β*_*i*_), *i* =1, 2, …, *g* be a non-singular odd characteristic. Then for any 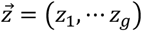 Fay’s identity is

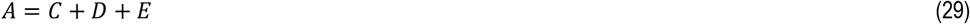

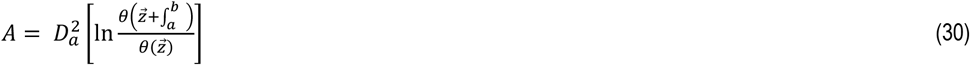

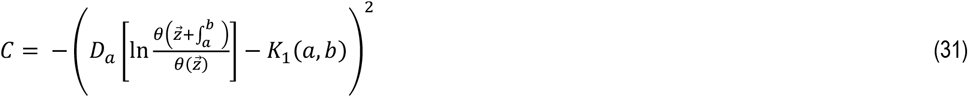

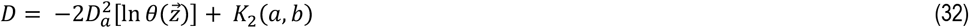

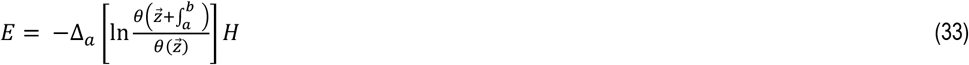

where 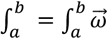 and the scalars *K*_*1*_ (*a, b*), *K*_*2*_ (*a, b*) are given by

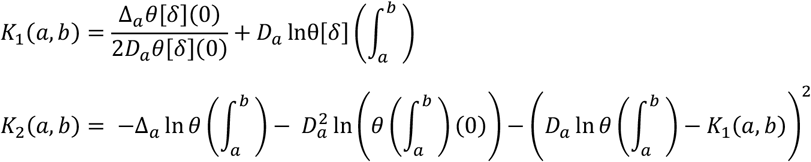

where *D*_*a*_ is the operator of directional derivate along the vector 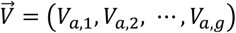 while *Δ*_*a*_ is the operator of directional derivative along the vector 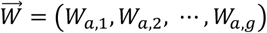 where these vectors describe the pinch distortion at a point of *Σ*_*g*_. Thus, the Fay’s identity in this form is for a specific pinch deformation. Substituting the multi-soliton solutions of the non-linear Schroedinger equation written down earlier it can be verified that they satisfy the degenerate Fay identity [31]. Thus, we have analytic expressions for these excitations in terms of theta functions with pinch-deformed variables. We are interested in the soliton limit of this solution.

The theta functions in the degenerate Fay identity have pinch deformed variable induced by the pinch deformation variables of the Riemann surface encoded in its one forms. The pinch deformations in greater detail can be written as *ω*_*j*_(*p*) = (*Va, j* (*p*) + *Wa, j* (*p*) *k*_*a*_ (*p*) + ⋯ )*dk*_*a*_(*p*), using the representation of the one form to parametrize the signal specific local pinch deformations. The range of the label *j* represents the number of pinch points that contribute to the soliton excitation.

### 7.2 The soliton limit

Soliton solutions emerge in the limit when the genus *g* signal producing surface, collapses to genus zero. The technical way of doing this has been previously discussed [31]. As a genus-zero surface is conformal to a Riemann sphere there is a map *w*(*p*) = *w* which maps a point *p* on the genus zero surface to a point *w* on the Riemann sphere. We will not use these solutions [31] as they have extra oscillation parts but a different one [34].

#### 7.2.1 Thalamic neuron spike data

##### METHODS

###### Viral inoculation

Young adult (p30) male mouse C57Bl/6J wild type was placed in deep anesthesia (isoflurane) and prepared for stereotaxic cranial inoculation. The dorsal part of the animal’s head was first shaved and then disinfected topically with ethanol 70%. Under the area a subcutaneous injection, (0.1ml) of amix of lidocaine 5mg/ml (Streuli) and Bupivacaine 1.25mg/ml (Sintetica). The mouse was placed in a dual stereotaxic frame (WPI) with drill and micropump. The animal received application of topic Carbomerum 980 2mg/ml (Bausch+Lomb) in both eyes and the body temperature was monitored and kept at 35°C (DC Temperature Controller, FHC), all through the surgery. The head skin was cleared with a vertical cut using a sterile scalpel blade. Once the area had been cleaned using sterile NaCl 0.9% serum solution (Braun), and the bleeding stopped, a small craniotomy was performed on the right parietal section of the skull to provide entry point to the inoculation. With an airtight precision syringe (10μl Hamilton, 1701 N ga26/51mm/pst3), slowly and gently moved the needle tip into the layer 5 of the right hemisphere S1 cortical area. Coordinates [57] to Bregma (in mm): M.L.: +1.5 ; A.P.: -0.5 ; D.V.: -1.4. There, a dose of the viral particle AAV9.CaMKIIa.ChR2(H134R).mCherry (Addgene), was inoculated (700nl: dilution 1:5) at a steady rate (100nl/min). Once it was finished the syringe stayed in position for at least 10 minutes before slowly and gently retreat and exit the tissue. The animal was then sutured, received a subcutaneous dose (0.3ml) of NaCl 0.9% serum (Braun) to help hydration and was placed in a recovery cage until was ready to return to his home cage. The animal was monitored through its recovery and had Ibuprofen 20mg/ml (Verfora) in the drinking water 24h prior and 72h post the surgery. All procedures to the animal were under the supervision and authorization of DGAV-affaires veterinaires du canton Vaud, Switzerland license VD3802, and according to EPFL veterinary regulations.

###### Coronal slice preparation

Adult (p56), male mouse C57Bl/6J wild type was placed in deep anesthesia (isoflurane) and then decapitated. The brain was quickly extracted and placed in cold oxygenated (O_2_ 95%; CO_2_ 5%) cutting extracellular solution (in mM: 213.0 sucrose, 2.5 KCl, 10.0 MgCl_2_, 1.25 NaH_2_PO_4_, 0.5 CaCl_2_, 25.0 glucose, 25.0 NaHCO_3_). In the same solution and with the assistance of a semiautomatic vibrating blade microtome (Leica VT1200S), acute coronal sections 300μm thick were obtained. The slices then were placed at room temperature in oxygenated (O2 95%; CO2 5%) normal extracellular solution (in mM: 125.0 NaCl, 2.5 KCl, 1.0 MgCl_2_, 1.25 NaH_2_PO_4_, 2.0 CaCl_2_, 25.0 glucose, 25.0 NaHCO_3_) for at least 1h to recover before recording. All chemicals were obtained from Sigma-Aldrich.

###### Whole-cell patch-clamp in current clamp mode recording

The slice (coordinates [57] to Bregma: A.P.: -2.30mm) was placed on a recording chamber at 33°C (Temperaturcontroller LandN) that was constantly perfused (peristaltic pump P-1, Amersham Biosciences) with oxygenated (O_2_ 95%; CO_2_ 5%) normal extracellular solution (in mM: 125.0 NaCl, 2.5 KCl, 1.0 MgCl_2_, 1.25 NaH_2_PO_4_, 2.0 CaCl_2_, 25.0 glucose, 25.0 NaHCO_3_). Neurons were located visually using an upright microscope (Olympus BX51WI) with IR-DIC optics and a digital camera (Prime95B Teledyne photometrics). Borosilicate capillaries with filament (Hilgenberg; 1403513) were pulled with a pipette puller (p-97, Sutter Instruments) into pipettes that were filled with intracellular solution (in mM:110.0 K-Gluconate, 10.0 KCl, 4.0 Mg-ATP, 10.0 Na-Phosphocreatine, 0.3 Na-GTP, 10.0 HEPES, 8.0 Biocytin; pH 7.3, 295mOsm). The intracellular electrode (Ag/AgCl) presented a resistance between 5 to 10MΩ in the bath. All chemicals were obtained from Sigma-Aldrich. A neuron from the Po thalamic nucleus was recorded in whole cell, current clamp, conditions using Axopatch 200B amplifiers (Axon Instrument). Data acquisition was performed via ITC-18, connected to a Windows based PC (Dell), running a custom-made routine in Igor Pro (V 9.0, Wavemetrics). The voltage signal was sample at rates between 5kHz, and filtered with a 2kHz Bessel filter. The injected current was to obtain -70mV at steady state, previous to stimulation. The response to opto-stimulation was recorded under these conditions. Opto-stimulation coupled to the microscope, a CoolLED pT-100, white light lamp controlled through digital output by a custom-made routine in Igor Pro (V 9.0, Wavemetrics), at 100% intensity, delivered light flashes through a filter cube (Olympus U-MNBV2, 475nm), and an objective (LUMPlanFI/IR; 40*x*/0.80*W*;∞/0) onto the recorded neuron and surroundings. The protocol was of 8 repetitions, duration 3ms, 20Hz, and 1 repetition 1s after first. Trace presented (Fig. 3) is one of these repetitions

#### 7.2.2 Memory structure

The non-transient helical spin structure formed by surface spins with the transient magnetic field generated by moving soliton pulses is determined by the quantum time evolution operator *U*(*t*) defined by the Hamiltonian *H*

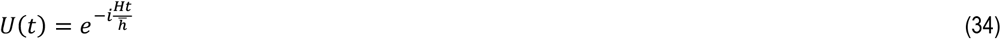

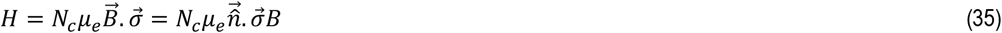

The Hamiltonian operator *H* describes the quantum interaction of spin-half particles with an external magnetic field, where *σ*_*i*_, i = 1, 2, 3 are Pauli matrices that represent spin-half particles [39]. We have written 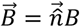 where 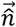 is the direction of the magnetic field 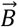 and *μ*_*p*_ = 2*μ*, where *μ* is the nuclear magnetron of a proton. The magnetic field interacts with *N*_*c*_ spins that are on a circle round the axon tube with centre *z*. It follows from the fact that the square of a Pauli matrix *σ* is equal to one, and 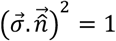, when 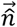 is a unit vector and that,

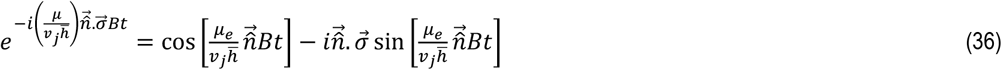

The time evolution of an initial spin *S j* (0) located at a point, is given by the equation, 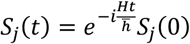. To study the spin distribution, parametrized by z, by replacing *t* by 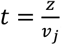, where *z* is a point in the centre of the axon tube and the spins are on the surface of the tube and *v*_*j*_ is the speed of the pulse. The spatial spin aligning evolution operator *U*(*z*), is given by,

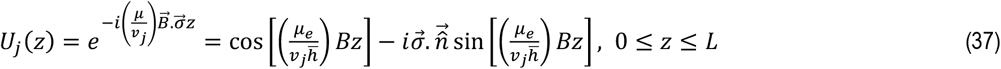

where the range of *z* defines the size of the memory structure. It is expected to have a closed topological loop structure. We write 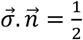, which is the aligned spin value projected out by the evolution operator and replace *μ*_*e*_ by 2*μ* the nuclear magnetron of the proton. Our static helical spin-magnet distribution, *S*_*j*_(*z*) is given by,

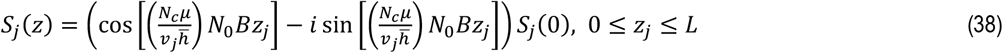

It is a helical chain of surface proton spins, where *S*_*j*_(0) represents the aligned spin at *z* = 0. It is created by the transient magnetic field pulses of a single soliton pulse.

### 7.3 Memory excitation frequency

The quantum Hamiltonian for describing the interaction of spin-half particles with a magnetic field *H* leads to a simple formula for the memory excitation frequency of each spin-half particle simply by writing the eigenvalue of the Hamiltonian *H* as 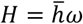. Thus,

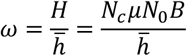

where 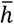 is Planck’s constant, *μ* the magnetic moment of a proton, *N*_*c*_ the number of spins on a circle of axon radius at a given point of the axon, and *N*_0_ is the average charge carried by a single pulse. The value of this frequency will depend on the values of *N*_*c*_, *N*_0_, *B*. We can estimate 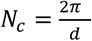, where *r* is the axon radius and *d* is the average separation distance between protons on the myelin sheath. Setting *r* ≈ 10^−4^cm and *d* ≈ 10^−8^cm, gives *Nc* ≈ 10^5^, we estimated *b* ≈ 10^−9^gauss, and thus *ω* ≈ *N*_0_Hz. Thus, the frequency depends on the charge carried by a soliton pulse. It also depends on the profile of the magnetic field and hence is signal dependent. We have ignored the detailed nature of *B* and the interaction between spins in this estimate. Any refined calculation should lead to excitation frequency values close to this value. We will call this the primary memory excitation frequency.

We showed that due to the special topological properties of spin half particles, there will always be a natural secondary memory structure present due to the pairing of two neighbouring spins of opposite orientation located at two neighbouring helical sites with double excitation frequency.

## Footnotes

1 In the Appendix we sketch how to construct the Riemann surface associated with the hyperelliptic equation and state some important features.

2 We will take the spin-half particles to be protons for reasons discussed elsewhere [41]

3 The corresponding *L* value for aligning electron spins is also 10cm, using a separation distance between electrons of 10^-7^cm, *μ* ≈ 10^−20^.

4 The possibility that proton spin can store memory has been suggested [51] in Posner molecules

## References

[1] Kandel E. (2002), In Search of Memory: The emergence of a new science of the mind, Norton and Company Inc.

[2] O’Sullivan F.M., Ryan T.J. (2024), If engram are the answer, what is the question? Advances in Neurobiology, 38, 273–302. DOI: 10.1007/978-3-031-62983-9_15.

[3] Scott A. (2002) Neuroscience: A Mathematical Primer. Springer-Verlag New York, Inc.

[4] Koch C. (2004) Biophysics of Computation, Oxford University Press

[5] Sompolinsky H., (2014) Computational Neuroscience: beyond the local circuit. Current Opinions in Neurobiology, 25, 13–18 DOI: 10.1016/j.conb.2014.02.002

[6] Sompolinsky H., Cristani A., Sommers H.J. (1988), Chaos in Random Neural Networks, Physical Review Letters, 61, 259–262 DOI: 10.1103/PhysRevLett.61.259

[7] Bialek W., Rieke F., de Ruyter van Stevenick R.R., Warland D. (1991), Reading a neural code, Science, 252, 1854–1857 DOI: 10.1126/science.2063199

[8] Anderson S.L., Jackson A.D., Heimburg T. (2009), Towards a thermodynamic theory of nerve pulse propagation, Progress in Neurobiology, 88, 104–113 DOI: 10.1016/j.pneurobio.2009.03.002

[9] Mussel M., Schneider M.F. (2019), It sounds like an action potential: unification of electrical, chemical and mechanical aspects of acoustic pulses in lipids, Journal of the Royal Society, Interface, 16, 20180743 DOI: 10.1098/rsif.2018.0743

[10] Shrivastava S., Schneider M.F. (2014) Evidence for two-dimensional solitary sound waves in a lipid controlled interface and its implications for biological signalling, The Royal Society Interface, 11, 20140098 DOI: 10.1098/rsif.2014.0098

[11] Tonegawa S., Pignatelli M., Roy D.H., Ryan T.J. (2015), Memory engram storage and retrieval, Current Opinion in Neurobiology, 35, 101–109 DOI: 10.1016/j.conb.2015.07.009

[12] Queenan B.N., Ryan T.J., Gazzaniga M.S., Gallistel C.R. (2017), On the research of time past: The hunt for the substrate of memory, Annals of the New York Academy of Sciences, 1396, 108–125 DOI: 10.1111/nyas.13348

[13] Ortega-de San Luis C., Ryan T.J. (2022) Understanding the physical basis of memory: molecular mechanisms of the engram, Journal of Biological Chemistry, 298 101866 DOI: 10.1016/j.jbc.2022.101866

[14] Lopez M.R., Wasberg S.M.H., Gagliardi C.M., Normandin M.E., Muzzio I.A. (2024) Mystery of the memory engram: History, current knowledge and unanswered questions. Neuroscience and biobehavioural reviews 159, 105574 DOI: 10.1016/j.neubiorev.2024.105574

[15] Ryan T.J., Roy D.S., Pignatelli M., Arons A., Tonegawa S. (2015) Engrams cells retain memory under retrograde amnesia, Science, 348, 1007–1013 DOI: 10.1126/science.aaa5542

[16] Semon R.W., (1904) Die Mneme: als erhaltendes Prinzip im Wechsel des osrganischen Geschehens, Leipzeig: Engelmann

[17] Tansey J.T. (2019) Biochemistry: An Integrative Approach Vol 1 p 143, John Wiley and Sons

[18] Feinberg G., Sucher J. (1970) General Theory of van der Waals Interaction: A Model Independent Approach, Physical Review A, 2, 2395 DOI: 10.1103/PhysRevA.2.2395

[19] Purcell, E.M. (1977) Life at Low Reynold Numbers, American Journal of Physics, 45, 3–11 DOI: 10.1119/1.10903

[20] Salzer J.L. (1997) Clustering sodium channels at node of Ranvier: close encounters of the axon-glia kind, Neuron, 18, 843–6 DOI: 10.1016/s0896-6273(00)80323-2

[21] Jackson J.D. (1962) Classical Electrodynamics, Wiley and Sons Inc.

[22] Nash C., Sen S. (1983), Topology and Geometry for Physicists, Academic Press

[23] Munkres J.R. (2000) Topology, Chapter 12, Prentice Hall, International Edition

[24] Lappalainen J.K., Tschopp F.D., Prakhya S., McGill M., Nern A., Shinomiya K. Takemura S.Y., Gruntman E., Mackle J.H. Turaga S.C. (2024) Connectome-constrained network predicts neural activities across the fly visual system, Nature, 634, 1132–1140 DOI: 10.1038/s41586-024-07939-3

[25] Pospisil D., Aragon M.J., Dorkenwald S., Matsliah A., Sterling A.R., Sclegel P., Yu S.C., McKellar C.E., Costa M., Eichler K., Jefferis G.S.X.E., Murthy M., Pillow J.W. (2024) The fly connectome reveals a path to the effectome, Nature, 634, 201–209 DOI: 10.1038/s41586-024-07982-0

[26] Atiyah M.F. (1971) Riemann Surfaces and Spin Structures, Annales Scientifique de l’Ecole Normale Supérieure, Serie 4, 4, 47–62 DOI: 10.24033/asens.1205

[27] Mumford D. (1971) Theta characteristics of an algebraic curve, Annales Scientifique de l’Ecole Normale Supérieure, Serie 4, 4, 181–192 DOI: 10.24033/asens.1209

[28] Mumford D. (1983) Tata Lectures on Theta, Chapter 1 & 3. Birkhäuser Boston, MA

[29] Teleman C. (2003), Riemann Surfaces, https://math.berkeley.edu/~teleman/math/Riemann.pdf

[30] Arbarello E. (2002) Sketches of KdV, Contemporary Mathematics, 312, 9–70 DOI:10.1090/conm/312/05391

[31] Kalla C. (2012) Thesis, Fay’s Identity in the theory of Integrable Systems, Université de Bourgogne. English. https://theses.hal.science/tel-00622289/file/these_A_KALLA_Caroline_2011.pdf

[32] Riemann B (2004) Collected Papers Translated from the 1892 German edition by Baker R., Christenson C., Orde H., Kendrick Press, Heber City, UT. https://www.scribd.com/document/482235342/Riemann-B-Collected-papers-Kendrick-2004-pdf

[33] Guardia J. (2002) Jacobian nullwerte and algebraic equations, Journal of Algebra, 253, 112–132

[34] Previato E. (1985) Hyperelliptic Quasi-Periodic and Soliton Solutions of the nonlinear Schroedinger Equation, Duke Mathematical Journal, 52, 329–377 https://projecteuclid.org/journalArticle/Download?urlid=10.1215/S0012-7094-85-05218-4

[35] Fay J. (1973) Theta functions on Riemann surfaces, Lectures Notes in Mathematics, 352.

[36] Sen S., Raina A. (1988). Grassmanians, Multiplicative Ward Identities and Theta Function Identities, Physics Letters B, 203, 256–262 DOI: 10.1016/0370-2693(88)90548-5

[37] Kalla C. (2011) Breathers and generalized non-linear Schroedinger equations as degenerations of algebro-geometric solutions, Journal of Physics A: Mathematical and Theoretical, 44, 335210 DOI: 10.1088/1751-8113/44/33/335210

[38] Faisal A.A., White J.A., Laughlin S.B. (2005) Ion-channel noise places limitations on the miniaturization of brain wiring, Current. Biology, 15, 1143–9 DOI: 10.1016/j.cub.2005.05.056

[39] Landau L.D., Lifshitz E.M. (1977). Quantum Mechanics: Non-relativistic Theory, Pergamon Press.

[40] Ritz T., Adem S., Schulten K. (2000). A model for photoreceptor based magnetoreceptors in birds, Biophysical Journal, 78, 707–718 DOI: 10.1016/S0006-3495(00)76629-X

[41] Sen S., (2025) A Topological method of generating action potentials and EEG oscillations in a surface network, Royal Society Open Science, 12, 241977 DOI: 10.1098/rsos.241977

[42] Barry J.F., Turner M.J., Schloss J.M., Glenn D.R., Song Y., Lukin M.D. Park H., Walsworth R.L. (2016) Optical magnetic detection of single-neuron action potentials using quantum defects in diamond, PNAS, 113, 14133–14138 DOI: 10.1073/pnas.1601513113

[43] Sterratt D., Graham B., Gillies A., Einevoll G., Willshaw D. (2023) Principles of Computational Neuroscience, Cambridge University Press

[44] Landau L.D., Lifshitz E.M. (1980) Statistical Physics, Pergamon Press

[45] BioNumbers, https://bionumbers.hms.harvard.edu/search.aspx

[46] Wang L. (2012) Measurements and implications of the membrane dipole potential, Annual Review of Biochemistry, 81, 615–635 DOI: 10.1146/annurev-biochem-070110-123033

[47] El Hady A, Machta B. (2019) Mechanical Surface waves accompany action potential propagation, Nature Communications, 6, 6697 DOI: 10.1038/ncomms7697

[48] Glatte P., Buchmann S.J., Hijazi M.M. Illigens B.M.W., Siepmann T. (2019) Architecture of the Cutaneous Autonomic Nervous System, Frontiers in Neurology, 10, 970 DOI: 10.3389/fneur.2019.00970

[49] Uchida M., Onose Y., Matsui Y., Tokura Y. (2006) Real Space observation of helical spin order, Science, 311, 359–361 DOI: 10.1126/science.1120639

[50] Vedmedenko E.Y., Altwein D. (2014) Topologically Protected Magnetic Helix for all spin based applications, Physical Review Letters, 112, 017206 DOI: 10.1103/PhysRevLett.112.017206

[51] Fisher M.P.A. (2015) Quantum Cognition: The possibility of processing with nuclear spin in the brain, Annals of Physics, 362, 593–602 DOI: 10.1016/j.aop.2015.08.020

[52] Amit D.J. (1989) Modeling brain functions: The World of attractor neural networks, Cambridge University Press

[53] Cook B.J., Peterson A.D.H., Woldman W., Terry J.R. (2022) Neural Field Models: A mathematical overview and unifying framework, Mathematical Neuroscience and Applications, 2, 1–67 DOI: 10.46298/mna.7284

[54] Sejnowski T., Koch C., Churchland P.S. (1988) Computational Neuroscience, Science, 241, 1299–1306 DOI: 10.1126/science.3045969

[55] Harnack A. (1876) Ueber die Vieltheiligkeit der ebenen algebraischen Curven, Mathematische Annalen, 10, 189–198 DOI: 10.1007/BF01442458

[56] Kallel S., Sjerve D. (2010) Invariant Spin Structures on Riemann Surfaces, Annales de la Faculté des Sciences de Toulouse, 19, 457–477 DOI: 10.5802/afst.1251

[57] Paxinos G., Franklin K.B.J. (2019) The Mouse Brain in Stereotaxic Coordinates, Academic Press, Elseiver

